# Bridging the Gap in Cancer Cell Behavior Against Matrix Stiffening: Insights from a Trizonal Model

**DOI:** 10.1101/2023.12.02.569730

**Authors:** Mohammad E. Torki, Fan Liu, Rongguang Xu, Yunfeng Chen, Jeffery Fredberg, Zi Chen

## Abstract

The intricate interplay between actomyosin contractility and extracellular matrix (ECM) strain stiffening is pivotal in cancer invasion. Despite the admitted impact of such feedback, current models are inadequate in predicting the largely overlapping ranges of cell shapes and their corresponding motility levels at intermediate ranges of collagen density. To address this gap, we introduce a free energy-based, trizonal model for cell shape transition under ECM stiffening, which delineates two distinct and one overlapping motility zones entitled with their implications for cancer progression: a low-motility zone with minimal invasiveness, a high-motility zone indicative of significantly invasive cells, and a mesoregion which harbors cells at crossroads of both states. This model integrates critical factors influencing the bidirectional interaction between the cell and ECM, thereby offering a deeper grasp of cancer cell behavior. Our findings reveal that the combined effects of ECM strain stiffening and cellular contractility are key drivers of cell population heterogeneity and invasiveness. This model goes beyond existing paradigms by accurately determining the optimal cell elongation at matrix-driven steady-state equilibrium, factoring in collagen density, contractility density, stress polarization, membrane-cortical tension, and integrin dynamics through the lens of total free energy minimization. The model’s predictive capability is further validated against measured cell shapes from histological sections. Altogether, this research not only bridges a crucial knowledge gap, but also provides a robust computational framework for predicting and replicating cell shape transitions observed in human functional tissue assays, thereby enhancing our ability to understand and potentially combat cancer invasion.

**Significance:** ECM stiffening is crucial in prompting metastatic phenotypes, with the interaction between cell contractility and ECM stiffening heavily influenced by cell motility level and reflected in distinct cell shapes [1–3]. This research introduces a free-energy-based model that, based on sound physics, not only distinguishes among different cell populations by their motility levels, but also truly replicates the recently observed trizonal cell response to ECM stiffness. This predictive model, validated by experiment, bridges a critical gap in our understanding of cellular dynamics in cancer progression, offering profound insight into the physical concepts driving these complex interactions. Thereupon, this work provides a powerful computational tool, potentially leading to new strategies in diagnosing and treating cancer by targeting specific cell behavioral traits and interactions within the tumor microenvironment.

## 1 Introduction

Collective cell migration occurs in many biological processes such as embryogenesis, tumorigenesis and wound healing. In all such processes, cells sustain interactions among themselves as well as with their surrounding tissues, with their shapes subject to continuous evolution. Notably, chemo-mechanically interacting intracellular and extracellular cues within the cell-matrix microenvironment are known as primary origins of epithelial-to-mesenchymal (EMT) as well as endothelial-to-mesenchymal transition (EndMT) underlying metastatic cell migration [4–6]. Recent experiments implied that cancer cells exhibited distinct collective invasion behaviors with varying extracellular ma-trix (ECM) concentrations. In particular, spheroidal cells residing in high-concentration collagen hydrogels (*i*.*e*., 4mg/ml) migrated slowly and collectively into finger-like shapes whereas those in low-concentration hydrogels (*i*.*e*. 2mg/ml) migrated rapidly and individually. The observed cell invasion behavior recapitulates the solid-fluid (melting and sublimation) phase transitions [7].

In general, cytoskeletal tension, actomyosin contractility, cell stiffness and extracellular matrix (ECM) fiber realignment are tightly correlated in malignant phenotypes [8–12]. The outward reflection of this multifaceted correlation is measurable and observable, respectively, through cell elongation and/or protrusion as well as cell contractility evaluated via the normalized PercevalHR ratio [12–16]. ECM strain stiffening and trans-membrane adhesion between the actin cytoskeleton and ECM are two salient factors affecting the prevailing cell shape. The latter is provided prevalently by integrins and less commonly through other adhesion receptors such as cadherins, selectins, and immunoglobulin-like cell adhesion molecules (Ig-CAM) [17]. Integrins act through focal adhesions that anchor the cell to ECM and transmit traction forces generated through actomyosin contractility [18–22]. Depending on the composition and linkage between their subunits, cell motility and invasion under the influence of integrins can be immensely enhanced [23], mitigated [24] or both [25–29]. Tolerances aside, high integrin expressions have a demonstrated capability of mediating multi-fold increase in cell contractility (reported for *α*5*β*1 for instance [22, 26–29]). The impact of ECM stiffness, however, is still in premature exploration. Indeed, a full-spectrum effect of ECM stiffness and its relevant parameters such as collagen density has remained elusive until recent years, thereby making some subtle controversy within the relevant literature. Earlier work suggest elevation of cell invasion upon observable increase in ECM stiffness in conjunction with matrix fiber alignment and increased collagen density [30–34]^1^. More recent findings, however, are indicative of multiphasic cell invasiveness in response to increasing collagen matrix stiffness [38–40], with maximum motility realized in an intermediate range of ECM stiffness ensued by remarkable reduction of invasion in highly stiff collagens. An analogous contrast is discernible in aged vs. young skin fibroblasts. As such, realignment of aged fibroblasts in stiff matrices promotes cell invasion whereas young fibroblasts compose a dense, isotropic matrix consistent with the so-called *basket weave* pattern [39, 41–43].

The highly anisotropic nature of metastasis is by and large rooted in matrix fiber realignment [44–47] or stress polarization [16, 48–50]. While the former is self-evident, the latter had been overlooked until quite recent times and the understanding of it still remains far from being adequate. The observed two-way feedback between cell protrusion and matrix alignment gives rise to a self-reinforcing mechanism involving interactive extension and retraction of cell protrusions [16, 50]. Accordingly, pre-aligned matrix fibers trigger alignment within cell protrusions which, in turn, leads to collagen fiber realignment. Realigned collagen fibers would ultimately stabilize the directions of protrusions by retracting their extension in lateral directions. Correspondingly, actomyosin contractility is primarily promoted by stress polarization, which is conventionally represented by directed springs in mathematical models. The “Clutch Model”, for instance, represents polarization with equivalent springs directed along cell protrusions [51, 52]. A distinct line of models were recently advanced to reflect the polarization effect in terms of the first principal stress (*σ*_1_) [16, 49, 50]. While well-motivated and partly consistent with physics, these models bare limited predictive capability from various perspectives. Organically, such models cannot capture the cell elongation/aspect ratio in correlation to matrix stiffness but defectively, *i*.*e*. through an abrupt (first-order) transition from spherical to elongated shapes. Apart from being non-physical by nature, such transition cannot capture the salient feature in recently observed cell-shape distributions at varying ECM stiffness. Zanotelli *et al*. [53] recently reported a representative case study wherein cell shapes vary from round to dramatically elongated over a large overlapping interval of collagen densities. This warrants an impending need to a new class of constitutive models that can predict two extreme (round vs. elongated) cell shapes in correlation with ECM stiffness. Section 1 of the Supplemental Material lays out, prior to derivation of the present model, a stepwise scheme for an extended formulation of models proposed in [16, 49, 50]. Corresponding plots of predicted cell body aspect ratio vs. ECM stiffness would then reveal the underlying shortcoming of such models and the warranted demand for the multizone model proposed herein. Hence, we hereby introduce a trizonal nonlinear constitutive model, similar in behavior to that proposed by Hannezo *et al*. [3] for epithelial cells, for migration of mesenchymal cells inside 3D matrix environments. The trizonal nature of the model yields three zones of contractility and optimal cell shape vs. ECM stiffness: (*i*) soft ECM zone characterized with low contractility and nearly spherical predominant cell shape, the two appearing insensitive to ECM stiffness; (*ii*) medium-stiffness ECM zone overlapping two possible solutions, the lower branch predicting nearly-spherical cell shape and low contractility, and the upper branch predicting elongated (*i*.*e*. invasive) cell shape and high contractility; (*iii*) high-stiffness ECM zone characterized with high contractility and cell elongation. The lower end to the intermediate stiffness zone would deliver an estimate of a crit-ical stiffness corresponding to intercellular de-adhesion and maximal cell invasiveness, later defined in Eq. (5). Numerical implementation of the model solidifies into a minimized total cell free energy with respect to strain and contractility as well as cell body aspect ratio. The former minimization is carried out a’ priori, expressing stress and contractility vs. strains whereas the latter minimization is executed a’ posteriori to predict the optimal cell shape primarily in terms of ECM stiffness. The distinct and combined effects of collagen stiffening, actomyosin and integrin inhibition are accordingly explored, with multifold applications: from predicting cancer-induced tissue impairment and cachexia, inflammatory and oxidative muscle injury to assessment analysis of therapeutic methods for reversal of such processes including Blebbistatin treatment of mesenchymal or epithelial cell cultures, HAPLN1 treatment of aged fibroblasts, myosin light-chain kinase inhibition, *etc*.

## 2 Model Derivation

### 2.1 STRESS-INDUCED SIGNALING DRIVES CELL CONTRACTILITY, ATP CONSUMPTION

In a sufficiently stiff extracellular matrix (ECM), intracellular contraction generates tensile forces in the matrix, significant enough to generate feedback with the cell through stress-sensitive signaling pathways (*e*.*g*., Rho-ROCK, Ca^2+^) [38, 54, 55]. Thereupon, continued binding of myosin motors and increasing contractile forces (through ATP consumption) would be promoted (Fig. 1a). By myosin motors undergoing the binding-force generation-unbinding cycle, cell contractility is maintained at a level proportional to the density of bound myosin. With ATP consumption required at each cycle, the energy released from ATP hydrolysis is converted into mechanical work and strain energy stored in the ECM as myosin motors contract the cytoskeleton until a steady state is realized. Thereupon, the entire energy from ATP consumption is converted to heat, with the dissipated contractile forces and ATP consumption varying with ECM stiffness. Stresses transmitted from the matrix through adhesions activate mechanosensitive signaling pathways, which upregulate the binding of myosin motors that leads to steady state. Each cycle of contractile force generation lasts within the order of milliseconds. In order for the cell to remain contractile over longer time scales, the cycles of binding, force generation and unbinding must operate continuously so ATP is continuously consumed [56–59].

**Figure 1:**
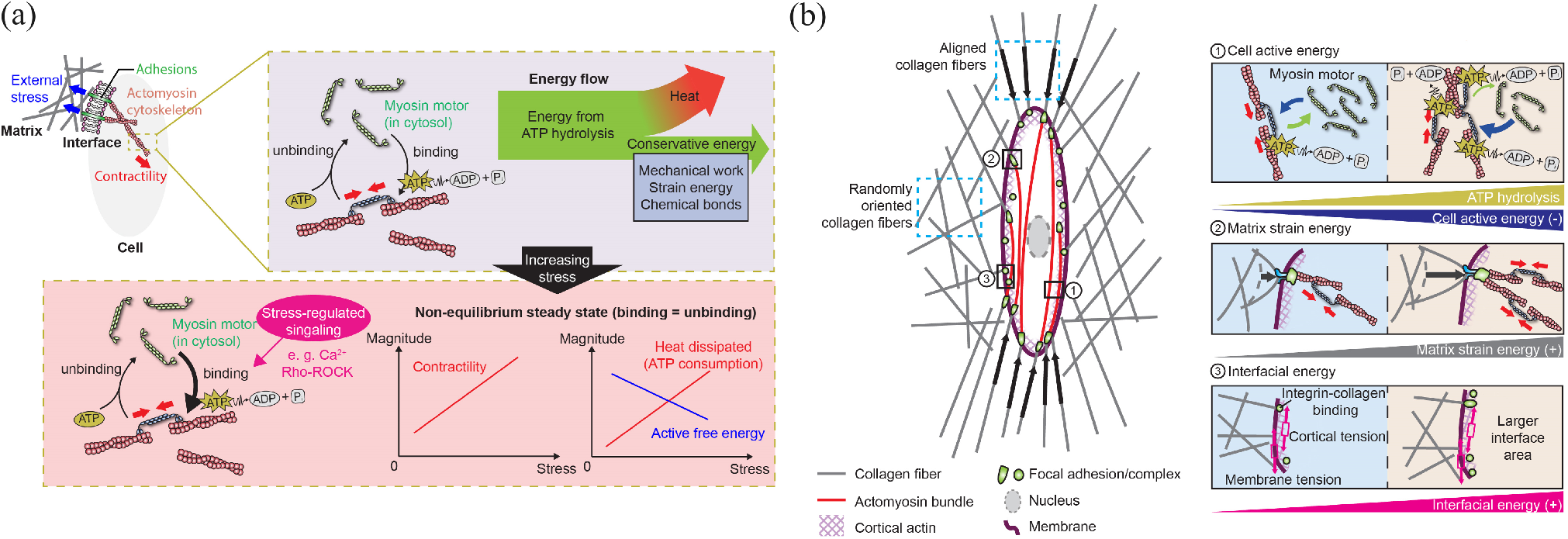
Chemo-mechanical representation of the coupling between stress-regulated signaling and myosin motor recruit-ment regulated by ECM strain stiffening. (a) Binding-force generation-unbinding myosin cycle until steady-state realization; (b) schematization of the total free energy contributed by the cell, matrix and interface. The competition among the corresponding energy components determines the optimal cell shape at steady state.

The complete cycle of binding and unbinding consists mainly of the binding/unbinding of individual myosin motors (in milliseconds) and phosphorylation of myosin motors (binding enabler), which is regulated by stress polarization through signaling pathways [56–59]. Accordingly, the active energy contributed by ATP hydrolysis normally manifests in the form of an exothermal process, hence appearing as a non-conservative (dissipative) term to be added to the cell’s overall free energy. That is, for a cell under one active stress-regulated signaling component:

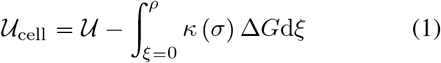

where *𝒰* represents the cell conservative energy constituted by mechanical, chemical and motility source terms (details are provided in Sec. 1 of the Supplemental Material), and the last term quantifies energy dissipation through ATP consumption. Existing linear models of cell durotaxis consider myosin motors to detach from the cytoskeleton immediately after the force generation cycle, thereby approximating *κ* as a linear function of stress. Such restriction, however, holds only under the circumstance of a large ATP resource, which proves overly restrictive under highly heterogeneous motor densities. With details saved for the Supplemental Material, the following expression for *κ* captures the highly anisotropic, multi-motile nature of signaling pathways between cell and ECM:

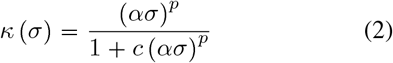

with *p, c* being non-dimensional parameters. Note that *p* ⩾ 1 and 0 ⩽ *c* ⩽ 1, whereby the linear feedback model would be retrieved with *p* = 1 and *c* = 0.

Next, due to the stress-dependent ATP hydrolysis being an exothermal process lowering the total free energy, the optimal cell shape is predicted by maximization of ATP consumption, hence minimizing the total free energy. The sequel expounds such process.

### 2.2 GENERALIZED STEADY-STATE CONDITION

The generalized steady-state condition can be attained through a minimization process. Accordingly, the time rates of contractile strains and forces (alias *contractility*) must vanish within the cell. To this end, the complete time derivative of total free energy can be expanded in terms of principal strain and contractility components as [33]

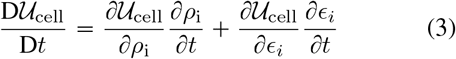

which, during steady state, entails that the conditions ∂*𝒰*_cell_/∂*ρ*_i_ 0 and ∂*𝒰*_cell_ /∂*ϵ*_*i*_ 0 be satisfied simultaneously.

Following the mathematical details presented in Sec. 1.2 of the Supplemental Material, the following nonlinear equation system will be derived in terms of the principal stresses *σ*_i_ and strains *ϵ*_*i*_:

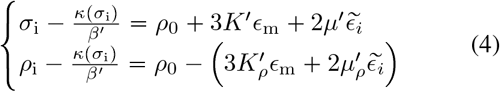

Where 3*K* ′ = 3*K* −1/*β*, 2*µ* ′ = 2*µ* −1*β* 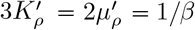 define the effective bulk and shear moduli and their counterparts for motor density, respectively. Moreover, *β*^′^ = *β/ ΔG*, and *κ* was defined in (2) as function of principal stresses. For validity of similitude among various parametric studies, model parameters are identified in terms of dimensionless parameters ς = *ρ*0/*E*, ℬ ≡ *βE* and Λ ≡ *α*/ *β*.

Provided the model parameters lie within their allowable ranges, Eq. (4) yields two possible solutions: one with limited contractility and corresponding polarization stress, and another with high contractility and corresponding polarization stress (see Sec. 2.2.1 of the Supplemental Material for the mathematical expose’ on the trizonal solution). The allowable ranges of model parameters (*α, β*), deliverable from the allowable (ℬ, Λ) diad, and tuning parameters (*p, c*) follow a rigorous asymptotic analysis over the special condition of the 1D domain. Section 3.1 of the Supplemental Material provides a self-contained elaboration on this matter.

Having obtained their principal components, the corresponding tensorial representation of stress and contractility can be determined from the polar decomposition theory,*i*.*e*. ***σ*** *=* σ_i_ ***n***^(i)^ ⊗***n***^(i)^ and *ρ = ρ*_i_ ***n***^(i)^ ⊗*n*^(i)^, where ***n***^(i)^ denote the principal directions, and summation is implied over *i*. For the sake of more clarity of model predictions as well as its parametric range, consideration of the one-dimensional cell-ECM subspace would be worthwhile. As such, the cell is ideally connected to a micropost (with its stiffness denoted with *K*_P_) or assumed to havea frozen spherical shape while being connected to a matrix with infinite radius. In order for better consistency with the general case, the post stiffness is conceived in terms of an equivalent matrix bulk modulus, whose equivalent Young’s modulus is identified in proportion to that of the cells as *E*_*M*_ *= ηE*, hence letting *K*_p_ = *E*_*M*_ */*3(1 −2 *µ*_M_.

## 3 Results

Figure 2 shows example plots of 1D trizonal solution (here represented by normalized stress) as well as an example case of post-processed optimum cell aspect ratios upon general 3D boundary-value implementation of the model at varying ECM stiffness ratios *η* (explanations to be seen in the sequel). The former reveals the effect of modeling tuning parameters *p* and *c*, noting that the linear model is retrievable by letting (*p, c*) = (1, 0). Increasing exponent *p* would chiefly decrease the upper-branch motility and contractile force level while increasing the unbinding-to-binding rate ratio *c* would would shrink the stiffness overlapping regime conducive to two possible cell shapes. Hence, equal binding and unbinding rates (*c =* 1) yields a neutral state with no or little distinction between lower and upper-branch motility levels and cell shapes. Altogether, tuning parameters best commensurate with physics of cell shape transition should lie within the ranges of 4 ⩽ *p* ⩽ 10 and 0.3 ⩽*c* <0.6.

**Figure 2:**
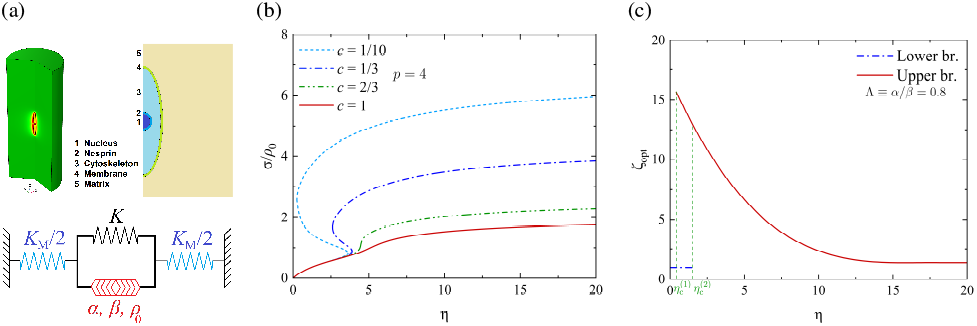
Trizonal model implementation. (a) Constituents of the 3D and 1D computational domains; (b) 1D trizonal solution for normalized stress vs. equivalent stiffness ratio *η* _ *E*_p_/*E* at various tuning parameters of *c* adopting (nearly) zero to 1, and *p* = 4; (b) representative example of predicted optimum cell aspect ratio from the lowly motile (lower branch) and highly motile (upper branch) solutions. Cell parameters are identified by *E* = 1.5 kPa, *ν* = 0.27, (*ς* _ *ρ*_0_/*E, B* _ *βE*, Λ _ *α*/*β*) = (0.2, 4.5, 0.8).

A significant corollary to the trizonal solution is the lower and upper bounds to the intermediate overlapping region, here denoted with 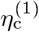 and 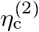. The lower bound physically delivers a matrix stiffness that corresponds to cell-ECM de-adhesion and initiation of maximal invasiveness. Upon detailed description provided in Sec. 2.2.1 of the Supplemental Material, 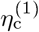 and 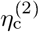 are expressible as

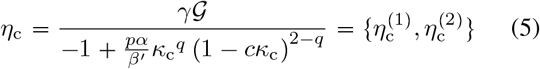

where 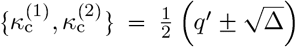, with 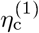 and 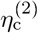 deliverable from 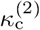 and 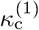, respectively. Here, 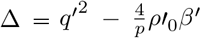, with *q* = 1 −1/*p* and 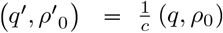. Further, *γ* and 𝒢 = *g* −1;*B* are problemdependent dimensionless parameters, such that 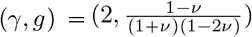 for a cell connected to microposts (provided the post stiffness is expressed in equivalent terms, as formerly noted), and (*γ, g)= (*1+ *ν*, 1 −2*ν)* for a cell connected to infinite ECM. The rest of parameters were defined in advance.

### 3.1 OPTIMUM CELL BODY ASPECT RATIO AT STEADY STATE

Elongation of a cell under the effect of cell-ECM mechanosensitive signalling pathways (*e*.*g*., Ca^2+^, Rho-ROCK) imparts significant alteration in the energetics of the entire domain. First, concentration of stresses near the elongated pole elevates stress upregulation of polarized signaling pathways thereby further recruiting myosin motors and formation of aligned contracted actin filaments, as indicated by increased first principal stress (*σ*_1_) and contractility (*ρ*_1_). Existing experimental results also demonstrate significantly higher actomyosin recruitment along with stronger bridging of focal adhesions by stress fibers in elongated fibroblasts on sufficiently stiff substrates [33, 35–37, 60]. Such increased actomyosin activity can be maintained by higher ATP consumption in the cell, reflected in reduction of the cell active energy 𝒰_cell_ upon cell elongation. In regards to the matrix, the significant deformation conveyed to the matrix at the cell poles would increase the strain energy stored in the matrix *𝒰*_M_. Such increase would reflect as an energetic penalty that prevents cells from taking on infinite aspect ratios by reducing the level of energy consumption in the combined cell-ECM domain. Finally, the interfacial energy is majorly reflected in membrane-cortical and integrin binding, and minorly attributed to membrane strain energy (see Sec. 2 of the Supplemental Material). Both components would increase with cell elongation, the former being in direct proportion to the cell outer surface area that increases with the cell body aspect ratio, and the latter due to elevated contractility stretching the adhesive bonds between the cell and ECM. Altogether, the competition between decreasing cell-ECM and increasing interfacial energy components leads to a non-monotonic profile of the total free energy *𝒰*_tot_ with respect to the cell body aspect ratio ζ (later observable in Fig. 3). Therefore, evaluation of the optimum cell body aspect ratio at a given steady state solidifies into a secondary (post-processed) stage that follows the evaluation of total free energy in terms of the cell body aspect ratio and spotting the aspect ratio that corresponds to the minimum energy.

**Figure 3:**
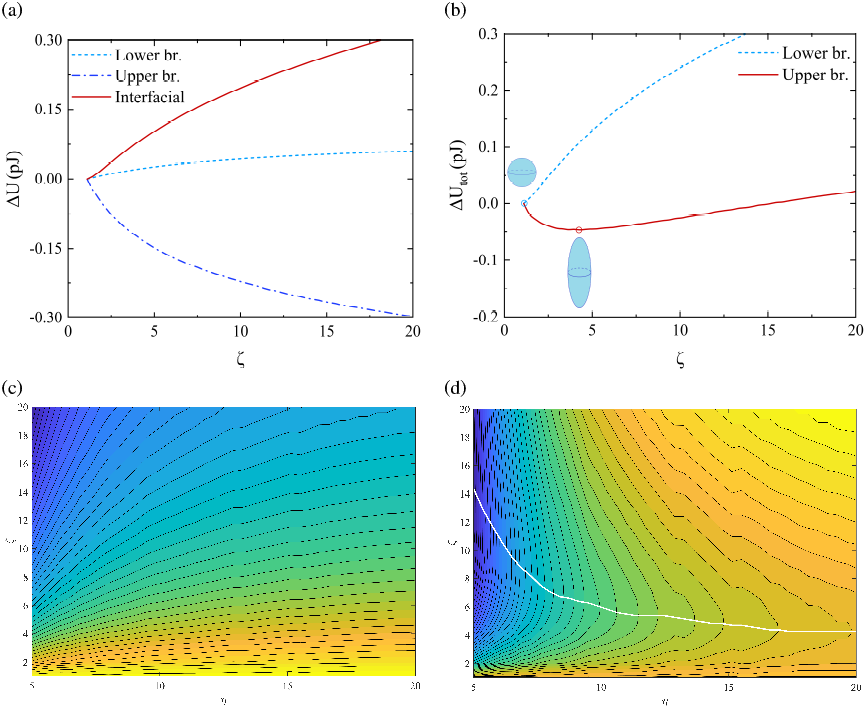
Free energy-based stiffness-regulated optimal cell shape at equilibrium state. (a) Variation of cell–ECM and interfacial free energy components (in reference to the spherical cell) vs. cell body aspect ratioζ at sufficiently large ECM stiffness (*η* _ *E*_M_/*E* = 5); (b) total free energy difference from superposition of cell–ECM and interfacial contributions; (c,d) lower and upper branch iso-energy contour lines, with their canyons (connected via white lines) denoting optimum aspect ratios (lower-branch canyons are realized entirely atζ = 1). Cell parameters and interfacial contribution are considered *E* = 1.5 kPa, *ν* = 0.27, (*ς, ℬ*, Λ) = (0.2, 5, 0.7) and (*γ*_0_, *γ*_1_) = (0.2, 0.001) mN/m, *δ*_ad_ = 1, respectively. Model tuning parameters are also fixed at (*p, c*) = (6, 1/3).

Note that, due to its limited ATP consumption and contractility, the lower branch of the constitutive model can only predict (nearly) spherical cell shapes. Hence, *𝒰*_tot_ corresponding to the lower branch is normally a monotonically increasing function of cell body aspect ratio, thus its optimum realized at ζ = 1. In the interest of better insight, more plots of ζ_opt_ vs. *η* (without energy contours) are shown and discussed in Sec. 2.2.2 of the Supplemental Material.

### 3.2 EXPERIMENTAL VALIDATION

The model’s parameteric predictability was assessed through a two-step machine-learning algorithm against cell-shape datasets from histological sections. They were subjected to the following reagents to the culture media of human HT1080 fibrosarcoma cells cultured in Dulbecco’s modified Eagle medium, and subsequently embedded in low-density (1.66 mg/mL) and mid-density (3 mg/mL) bovine collagens [61] (see Appendix A for details): (*i*) Blebbistatin (20 *µ*M, Sigma, 203391) and DMSO (0.1%, vehicle control, Sigma Aldrich) for actomyosin inhibition and its control culture, respectively; (*ii*) clone 4B4 mouse monoclonal anti-human CD29 (5 *µ*g/mL, Beckman Coulter, 6603113) and mouse IgG1 isotype control culture (5 *µ*g/mL, BD Bioscience, 557273); (*iii*) combination of Blebbistatin and clone 4B4 compared with the control culture with DMSO and IgG1 isotypes combined. Cell body aspect ratios (major over minor axis ratio) were approximated from mid-cellular cross sections exploiting an ellipsoid approximation plugin in Fiji/ImageJ (version 1.51). The matrix stiffness is hereby expressed in direct terms of the collagen density in reference to the evaluations using the *Atomic Force Microscopy* (AFM) [62–65]. Accordingly, the low and intermediate bovine collagen densities correspond to stiffness ratios approximated as *η* 1.75 and *η* 3.5, respectively.

The in-vitro histograms shown in Fig. 4 are the outcomes of a hybrid machine learning algorithm constructed from two subroutines. Thereupon, the best parameter is firstly adopted such that the least squared difference is acquired between the model predictions and the main data peaks on the histogram of interest. Secondly, the percentage distribution is simulated by application of a replicated multimodal probability distribution subroutine. Note that the nearly spherical cell body aspect ratios (1 ⩽ζ <1.5) were predicted by the model’s lower branch and higher ratios were evaluated using the upper branch (see Sec. 3.2 of the Supplemental Material for further information).

**Figure 4:**
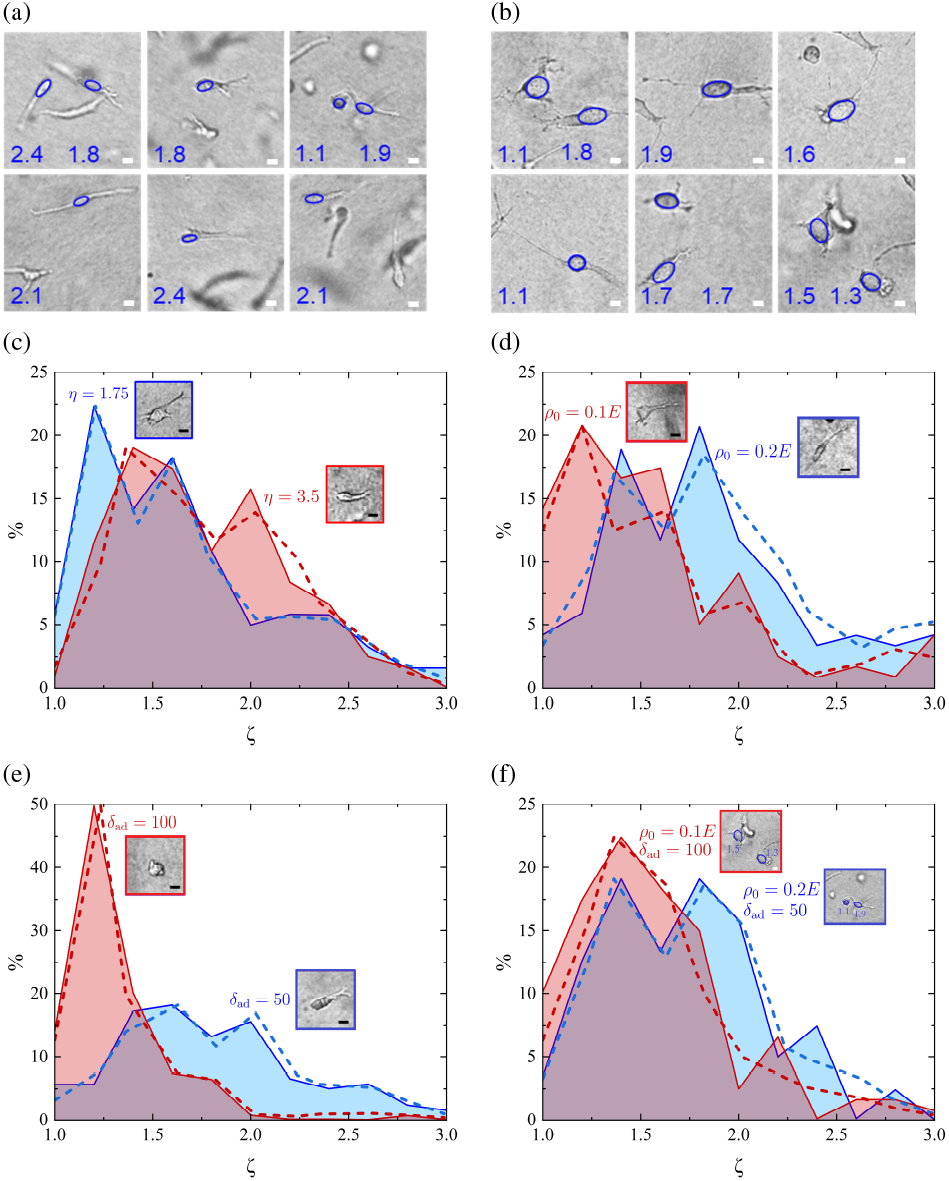
Representative cell morphologies showcasing phenotype heterogeneity. Morphologies of HT1080 cells migrating for 20 hours in 3 mg/ml bovine collagen with and without dual interference of Blebbistatin and 4B4 treatment: (a) example control assay treated with DMSO (0.1%, vehicle control, Sigma Aldrich) and mouse IgG1 isotype (5 *µ*g/mL, BD Bioscience, 557273) controls; (b) example assay treated with Blebbistatin (20 *µ*M, Sigma, 203391) and mouse monoclonal anti-human CD29 (clone 4B4, 5 *µ*g/mL, Beckman Coulter, 6603113). Ellipses indicate cell bodies with their respective aspect ratios (scale bars are 10 *µ*m); (c–f) predicted vs. experimental histograms of cell body aspect ratios: (c) for cells cultured in 1.6 and 3.0 mg/mL collagens, (d) for cells treated with Blebbistatin compared with the DMSO control histogram, (e) for cells treated with 4B4 mAb compared with the IgG1 isotype control histogram, (f) for cells treated with combined Blebbistatin–4B4 mAb, compared with the DMSO–IgG1 control histogram.

The first effect explored pertains to matrix stiffening through varying collagen density (Fig. 4c). Having cultured isolated melanoma cell spheroids in bovine collagen, cells were separated from the melanoma and invaded the matrix [38]. At low collagen concentrations (*e*.*g*. 0.1 mg/mL), cells invading at the periphery of melanoma had a spherical dominant morphology. With first-level increase in collagen density (into 0.5 and 1.0 mg/mL), the cells became clearly elongated. Upon second-level increase, cells became more elongated with some common aspect ratios with the former plus other spindle-like cells protruding from melanoma. Note that, since cells embedded in collagen densities clearly below 1.6 mg/mL were found to be insensitive to matrix stiffening, the model was tuned so as to predict *η =* 1.75 slightly above the minimum critical limit 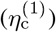 and *η =* 3.5 slightly above the maximum critical limit 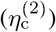, using tuning parameters of *(p, c*) = (10, 0.5). Accordingly, increasing collagen density from 1.6 to 3 mg/mL would maintain a considerable overlapping range of aspect ratios (represented by the common area between histograms) and simultaneously transfer an additional percentage to the highly motile (upper) branch (the pink area protruded around ζ = 2).

Next, actomyosin inhibition was investigated through treatment of HT1080 cells with Blebbistatin (Fig. 4d). While various local effects (including individual cell properties and local collagen fiber microstructures) can alter the measurable cell body aspect ratios, an overall decreasing effect is observable on cell elongation under such treatment, in comparison to the control culture treated with DMSO. It is thus conspicuous from the numerical histograms that decreasing the contractile density *ρ*_0_ by the appropriate range is directly conducive to this effect. Note that an indirect path to numerical replication of such effect is through increasing the polarization parameter or its normalized equivalent Λ while being maintained above its lower bound Λ_inf_. The reason lies intuitively in the direct correlation between the contractile density and mechanosensitive feedback triggered by stress polarization at a given cell steady state. Such effect is well commensurate with previous findings illustrating the effect of Y-27632 as a Rho-ROCK inhibitor [55]. Reduction of cell elongation thereunder further corroborates the essence of increased actomyosin activity for cell elongation and polarization. Moreover, the 4B4 monoclonal antibody (mAb) treatment would impair collagen binding through *β*1 integrins [66–68]. Through corresponding reduction of adhesion density *δ*_ad_ (Fig. 4e), cell elongation and polarization would be mitigated due to impairment of cell-matrix interaction, which downregulates mechanosensitive feedback, actomyosin activity and ATP consumption. Such effect is further commensurate with previous findings, *e*.*g*. reported by Castelló-Cros *et al*. who observed the same reduction effect on MDA-MB-231 cells under mAb13 treatment [68]. Note also that the effect of integrin impairment would render the aspect ration distribution unimodal, with its median driven towards unity, here around 1.2.

To be importantly noted, just as mere variation of collagen density (Fig. 4c) would prompt higher dispersity of the aspect ratio for its values up to 2, so too would both simultaneous treatments (Blebbistatin–4B4 and DMSO– IgG1) exhibited in Figs. 4(d,e). Existence of large diversity within cell shapes inside this range warrants the demand for a trizonal model that can easily output the largely distinct cell shapes at every given percentage.

To sum up, histological shape distribution patterns can be successfully replicated and shown to be consistent with predictions from the proposed analytical model (supplemented with machine learning algorithms), which emanate from the three fundamental regimes of cell shape transition predicted in its basic form.

## 4 Discussion

Insofar as steady-state cell shape is directly relevant to the competition amongst cell free energy, cell-ECM interfacial energy and ECM strain energy, the impact of pharmacological treatments on cell configuration best reflects in the energetics of both the cell and ECM. The myosin inhibiting trace of Blebbistatin in the foregoing Section, for instance, has been likewise observed from similar treatment on human bronchial epithelial cells (HBECs) [60]. Therein, Blebbistatin treatment would downregulate both actomyosin contractility and glycolysis through inhibiting myosin II accompanied by isolation of actin monomers and shrinkage of F-actin bundle lengths. The observed feedback among substrate stiffness, actomyosin activity and glycolytic rate lends further credibility to the introduced durotaxis model. Meanwhile, fibroblasts treated with Blebbistatin would turn considerably more dendritic and generate long, densely populated protrusions. Hence, obtaining an accurate estimate of the aspect ratio would entail fitting an ellipse to the innermost cell boundary [69]. As schematically shown in Fig. 5b, collagen fibers realign in the direction of stress-regulated polarization, providing contact guidance for formation of protrusions. Accordingly, an elongated cell tends to develop protrusions at both poles.

**Figure 5:**
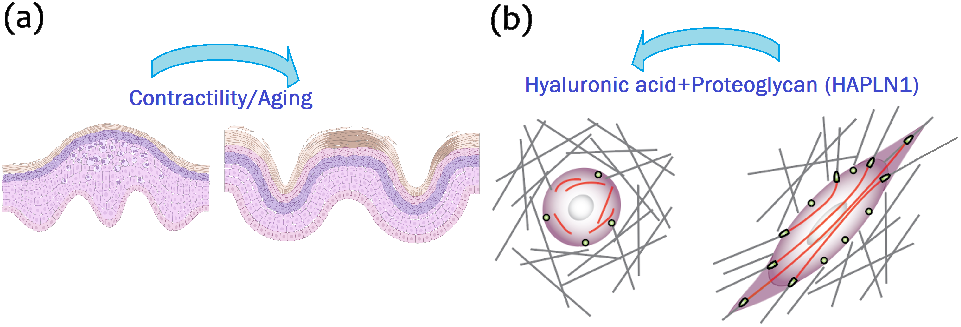
Extracellular strain stiffening and stiffness degradation can both instigate malignant phenotypes. (a) Schematic histological sections of young vs. aged human skin; (b) corresponding schematic cell morphology in young vs. aged skin, with elongation and ECM fiber-mediated protrusion prompted by interactive intracellular contractility, extracellular stiffening and fiber alignment.

A similar effect to that imparted by Blebbistatin treatment is achievable from hyaluronic and proteoglycan link protein (HAPLN1) in aged skin fibroblasts to revert realignment of fibroblasts in stiff matrices (Fig. 5) [39, 41–43]. In a recent endeavour by Kaur *et al*., such reconstruction was analyzed using the established in-vitro *cell-derived matrix* (CDM) technique to fabricate ECM and recapitulate aging-induced fiber realignment by mere means of fibroblasts [39]. They further extended the invitro data into an in-vivo model through assessment of ECM structure in young vs. aged C57BL/6 mouse skins. In both models, a remarkable difference was observable between young and aged fibroblasts in the sense that malignant cell migration would be promoted in dermal substrates whose stiffness had been degraded by age-related fibroblast realignment. Application of HAPLN1 would then reverse the dermal degradation mechanism whereby the so-called *basket weave* structure would be retrievable. The trizonal predicted cell elongation vs. ECM stiffness is true to the spirit of demoted cell contractility and elongation on overly stiff and HAPLN1-restructured substrates. This adds another layer of credibility to the proposed model. In more general sense, the demoted actomyosin activity and cell elongation inside overly stiff matrices is best reflected in the effect of reduced pore size with increasing collagen concentration. Thereupon, the confinement from collagen densification would preclude fibers from realignment, hence limiting cell shape transition. In particular, in an environment where matrix pores are much smaller compared to the cell volumes, cells must degrade and remodel the ECM network by secreting matrix metalloproteinase (MMP) before shape transition can occur. Therefore, in a non-degradable matrix or with deficient MMP secretion, cells would adapt to the shape of network pores [61, 70].

Apropos of the effect of cell-ECM adhesion, the effect induced by 4B4 antibody is similar to that mediated by *vinculin knockout* fibroblasts, which inhibit contractile force transmission from the cell on the surrounding matrix. Consequently, the cells would exhibit limited elongation and polarization [69]. By the same token allowing for predicting the effect of the 4B4 integrin inhibitor, the present model can very well quantify the sequestering effect of vinculin knockout fibroblasts. In fact, the formerly defined adhesion density *δ*_ad_ represents the density of collagen ligands connected to the cytoskeleton. Hence, cells would adopt less elongated configuration with fewer available ligands. Thus far, we have deemed no interdependence between interfacial parameters and the cell shape, hence rendering optimal cell shape function in terms of independently arbitrated membrane-cortical stiffness *γ*_0_ and adhesion density *δ*_ad_. While such approach is both qualitatively and quantitatively true to the spirit of cell shape transition, relevant studies allude to possible correlation at least between adhesion binding (mainly through integrins) and the instantaneous (unsteady) cell configuration [22]. Another line of prospective research would thus examine such possible correlation.

## Acknowledgements

Authors genuinely acknowledge support from the National Science Foundation under grant No’s 1953572, 1548571 and National Institute of Health under grant No. 6463801. Further acknowledgement goes to Prof. V. Shenoy, who provided constructive feedback on model predictions. Last but not least, experimental support from Prof. P. Friedl (Division of Cancer Medicine, MD Anderson Cancer Center), and technical support from Xingyu Chen (former PhD student and current Machine-Learning engineer) for inputting data for the linear model and providing sketches of cell–ECM interaction.

## Appendix

A Cell culture and collagen matrix

Human HT1080 fibrosarcoma cells (ACC315; DSMZ Braunschweig) were cultured in Dulbecco’s modified Eagle medium (DMEM; Sigma Aldrich), supplemented with 10% fetal bovine serum (FBS; Sigma Aldrich), penicillin (100 U/mL), streptomycin (100 mg/mL; PAA), and sodium pyruvate (1 mM; Thermo Fisher Scientific). Cell identity was verified by the Assay coded SNP–ID (from the Sequenom, MassArray System, Characterized Cell Line Core Facility, MD Anderson Cancer Center, Houston, TX), and freedom from contamination with mycoplasma was regularly verified using the MycoAlert Mycoplasma Detection Kit (Lonza). Cells from the sub-confluent culture were subsequently detached by trypsin-EDTA (2 mM, Sigma Aldrich) and embedded in low-density (1.66 mg/mL) and mid-density (3 mg/mL) bovine collagens (PureCol, Advanced Biomatrix) [61]. Following collagen polymerization (at 37°C for 20–30 minutes), single-cell cultures in the 3D collagen were submerged in the culture medium and maintained at 37°C. Finally, cell shape of individual cells were measured in 3D collagen cultures after 24 hours after the setting of single-cell spreading.

All antibodies and inhibitors used were purchased as azide-free or dialyzed (Slide-A-Lyzer mini dialysis device, 20 MWCO) compounds. Single-cell cultures in 3D collagens were incubated (37°C for 20h), and cell shapes were monitored by 3D brightfield microscopy (20X/0.30 HI Plan objective, DM IL LED microscope, Leica). Z-stacks were recorded, with one in-focus slice of each cell used for analysis. Cell body aspect ratios (major over minor axis ratio) were approximated from mid-cellular cross sections exploiting an ellipsoid approximation plugin in Fiji/ImageJ (version 1.51).

## Footnotes

1 Accordingly, the term "durotaxis" conventionally refers to migration of cells from a more compliant to a stiffer matrix [33, 35–37].

