## Supplemental Material for "Bridging the Gap in Cancer Cell Behavior Against Matrix Stiffening: Insights from a Trizonal Model"

#### 1 Formulation of Anisotropic Two-Way Feedback Model

The proof of concept for formulation of a complete cycle of actomyosin recruitment can be laid out with focus on 1D variation, with only one independent stress and contractility component. Denoting the binding and unbinding time rates by $J_{\text{on}} > 0$ and $J_{\text{off}} < 0$, the time rate of changing motor density, technically termed *contractility*, can be written as

$$\dot{\rho} = J_{\text{on}} + J_{\text{off}} \tag{S-1}$$

To ensure the positivity of energy dissipation rate, the following forms are adopted for $J_{\text{on}}$ and $J_{\text{off}}$:

$$J_{\text{on}} = -k_{\text{on}} \left( \frac{\partial U}{\partial \rho} - \Delta G \right) \quad , \quad J_{\text{off}} = -k_{\text{off}} \left( \frac{\partial U}{\partial \rho} \right) \tag{S-2}$$

with $k_{\text{on}} > 0$ and $k_{\text{off}} < 0$ being kinetic parameters for binding and unbinding rates, and $\Delta G$ is a scale factor representing energy realesed through ATP hydrolyzation [1,2]. The steady state entails $\dot{\rho} = 0$, hence writing

$$\frac{\partial U}{\partial \rho} = \kappa \Delta G \tag{S-3}$$

where $\kappa = \frac{k_{\text{on}}}{k_{\text{off}} + k_{\text{on}}}$.

The complete cycle of binding and unbinding consists mainly of the binding/unbinding of individual myosin motors (in milliseconds) and phosphorylation of myosin motors (binding enabler), which is regulated by stress through signaling pathways [3,4,5,6]. To capture the latter effect, $k_{\text{on}}$ should be set as a function of the external stress $\sigma$. In existing linear models of cell durotaxis, myosin motors are taken to detach from the cytoskeleton immediately after the force generation cycle, thereby giving rise to a significantly higher unbinding probability. Accordingly, $k_{\text{off}} >> k_{\text{on}}$ and $k_{\text{off}}$ is normally regarded constant. Such restriction, however, holds only under the circumstance of a large ATP resource, which proves overly restrictive under highly heterogeneous motor densities. Rather, highly motile cell shape transition under heterogeneous motility cannot be duly captured unless the binding rate is maintained above the unbinding rate, *i.e.* $k_{\text{on}} = (\alpha\sigma)^p \, k_{\text{off}}$, with $\alpha$ admitting a dimension of L$^2$/F. Such correlation captures the highly anisotropic nature of signaling pathways between cell and ECM. On the other hand, heterogeneity within

the cell motility level at a given quiescent contractility level, *i.e.* at fixed contractile density $\rho_0$, calls for consideration of a fraction of the steady-state unbinding rate, here denoted with a constant parameter such as $c$. Altogether, the following mathematical form proves true to both spirits:

$$\kappa\left(\sigma_i\right) = \frac{\left(\alpha\sigma_i\right)^p}{1 + c\left(\alpha\sigma_i\right)^p} \tag{S-4}$$

with $(p, c)$ being non-dimensional parameters. Note that $p \geqslant 1$ and $0 \leqslant c \leqslant 1$, whereby the linear feedback model would be retrieved with $p = 1$ and $c = 0$. By virtue of Eq. (S-4), increases in the principal stress $\sigma$ would facilitate the phosphorylation and creates a non-equilibrium steady state with increasing levels of cell contractility $\rho$ and ATP consumption. Accordingly, the active energy contributed by ATP hydrolysis must be added to the cell's conservative free energy. That is

$$\mathcal{U}_{\text{cell}} = \mathcal{U} - \int_{\xi=0}^{\rho} \kappa\left(\sigma\right)\Delta G \mathrm{d}\xi \tag{S-5}$$

where $\mathcal{U}$ represents the cell conservative energy constituted by mechanical, chemical and motility source terms (details are provided in Materials and Methods), and the last term quantifies energy dissipation through ATP consumption.

Upon recollection, the cell active free energy in its basic form is superposed by an active and a passive component from ATP consumption [7, 1, 8]:

$$\mathcal{U}_{\text{cell}} = \mathcal{U}_{\text{act}} + \mathcal{U}_{\text{pas}} \tag{S-6}$$

The active energy consists of the chemical energy associated with myosin recruitment (ATP hydrolysis) and the mechanical work from conversion of ATP hydrolysis to motor work [9]:

$$\mathcal{U}_{\text{act}} = \mathcal{U}_{\text{chem}} + \mathcal{U}_{\text{mot}} \tag{S-7}$$

where

$$\mathcal{U}_{\text{chem}} = \frac{\beta}{6}\left(\rho_{\text{kk}} - 3\rho_0\right)^2 + \frac{\beta}{2}\tilde{\rho}_{ij}\tilde{\rho}_{ij} \quad , \quad \mathcal{U}_{\text{mot}} = \rho_{\text{m}}\epsilon_{\text{kk}} + \tilde{\rho}_{ij}\tilde{\epsilon}_{ij}$$

with the $\tilde{(\cdot)}$ script denoting deviatoric components.

All the same, the cytoskeletal portion of the passive energy (excluding the energy stored in the nucleus and nesprin) reflects the mechanical energy stored therein upon being deformed by contractile forces. It can further be dissected into the stored strain energy minus the mechanical work done thereby. Each term can be resolved into two parts resulting from the mean and deviatoric strain components as [9]

$$\mathcal{U}_{\text{mech}} = \mathcal{U}_{\text{s}} + \mathcal{W} \tag{S-8}$$

where

$$\mathcal{U}_{\text{s}} = \frac{K}{2}\epsilon_{\text{kk}}^2 + \mu\tilde{\epsilon}_{ij}\tilde{\epsilon}_{ij} \quad , \quad \mathcal{W} = -\left(\int_0^{\epsilon_{\text{kk}}} \sigma_{\text{m}}\mathrm{d}e_{\text{kk}} + \int_0^{\tilde{\epsilon}_{ij}} \tilde{\sigma}_{ij}\mathrm{d}\tilde{e}_{ij}\right)$$

with $0 \leqslant e_{ij} \leqslant \epsilon_{ij}$, hence $0 \leqslant e_{\text{kk}} \leqslant \epsilon_{\text{kk}}$ and $0 \leqslant \tilde{e}_{ij} \leqslant \tilde{\epsilon}_{ij}$ denoting integral variables.

##### 1.1 Linear anisotropic model

Linearized approximation of the chemomechanical feedback is retrievable upon letting $(p, c) = (1, 0)$ in Eq. (S-4), the corresponding energy term is expressible in linear correspondence with stress components [9]. That is, in basic form:

$$\mathcal{U}_{\text{chem}} = \frac{\beta}{6}\left(\rho_{\text{kk}} - 3\rho_0\right)^2 + \frac{\beta}{2}\tilde{\rho}_{ij}\tilde{\rho}_{ij} - \left(\int_0^{\rho_{\text{kk}}} \alpha_{\text{v}}\sigma_{\text{m}}d\zeta_{\text{kk}} + \alpha_{\text{d}}\int_0^{\tilde{\rho}_{ij}} \tilde{\sigma}_{ij}d\tilde{\zeta}_{ij}\right) \tag{S-9}$$

which, with the additional term $\alpha_{\mathrm{a}}$ (after Shenoy and coworkers [10,11,12]), would extend to its anisotropic counterpart reading

$$\mathcal{U}_{\mathrm{chem}} = \frac{\beta}{6}\left(\rho_{\mathrm{kk}} - 3\rho_0\right)^2 + \frac{\beta}{2}\widetilde{\rho}_{ij}\widetilde{\rho}_{ij} - \left(\int_0^{\rho_{\mathrm{kk}}} \alpha_{\mathrm{v}}\sigma_{\mathrm{m}} d\zeta_{\mathrm{kk}} + \alpha_{\mathrm{d}}\int_0^{\widetilde{\rho}_{ij}} \widetilde{\sigma}_{ij} d\widetilde{\zeta}_{ij} + \int_0^{\rho_1} \alpha_{\mathrm{a}}\sigma_1 \mathrm{d}\,\rho_1\right) \tag{S-10}$$

where $\alpha_{\mathrm{m}}$, $\alpha_{\mathrm{d}}$ and $\alpha_{\mathrm{a}}$ are chemomechanical coupling parameters. The last one accounts for anisotropic stress polarization along the first principal stretch direction, hence conjugating with the first principal stress $\sigma_1$.

Recast in terms of principal variables, the total cytoskeletal portion of the total energy density is expanded as

$$\begin{aligned}\mathcal{U}_{\mathrm{cyt}} &= \frac{K}{2}\left(\epsilon_1 + \epsilon_2 + \epsilon_3\right)^2 + \mu\left(\widetilde{\epsilon_1^2} + \widetilde{\epsilon_2^2} + \widetilde{\epsilon_3^2}\right) - \frac{1}{6}\left(\sigma_1 + \sigma_2 + \sigma_3\right)\left(\epsilon_1 + \epsilon_2 + \epsilon_3\right) \\ &- \frac{1}{2}\left(\widetilde{\sigma_1}\widetilde{\epsilon_1} + \widetilde{\sigma_2}\widetilde{\epsilon_2} + \widetilde{\sigma_3}\widetilde{\epsilon_3}\right) + \frac{\beta}{6}\left(\rho_1 + \rho_2 + \rho_3 - 3\rho_0\right)^2 + \frac{\beta}{2}\left(\widetilde{\rho_1^2} + \widetilde{\rho_2^2} + \widetilde{\rho_3^2}\right) \\ &- \frac{\alpha_{\mathrm{v}}}{6}\left(\sigma_1 + \sigma_2 + \sigma_3\right)\left(\rho_1 + \rho_2 + \rho_3\right) - \frac{\alpha_{\mathrm{d}}}{2}\left(\widetilde{\sigma_1}\widetilde{\rho_1} + \widetilde{\sigma_2}\widetilde{\rho_2} + \widetilde{\sigma_3}\widetilde{\rho_3}\right) - \frac{\alpha_{\mathrm{a}}}{2}\sigma_1\rho_1 + \rho_1\epsilon_1 + \rho_2\epsilon_2 + \rho_3\epsilon_3\end{aligned} \tag{S-11}$$

Using the easily justifiable idendity

$$\frac{\partial\widetilde{x}_i}{\partial x_j} = \frac{\partial}{\partial x_j}\left(x_i - \frac{x_{\mathrm{kk}}}{3}\right) = \delta_{ij} - \frac{1}{3} \tag{S-12}$$

with $x$ representing $\sigma$, $\epsilon$ and $\rho$, one can write

$$\begin{aligned}\frac{\partial\mathcal{U}_{\mathrm{cyt}}}{\partial\rho_1} &= \frac{\beta}{3}\left(\rho_1 + \rho_2 + \rho_3 - 3\rho_0\right) + \frac{\beta}{3}\left(2\widetilde{\rho}_1 - \widetilde{\rho}_2 - \widetilde{\rho}_3\right) - \frac{\alpha_{\mathrm{v}}}{3}\left(\sigma_1 + \sigma_2 + \sigma_3\right) - \frac{\alpha_{\mathrm{d}}}{3}\left(2\widetilde{\sigma_1} - \widetilde{\sigma_2} - \widetilde{\sigma_3}\right) - \alpha_{\mathrm{a}}\sigma_1 + \epsilon_1 \\ \frac{\partial\mathcal{U}_{\mathrm{cyt}}}{\partial\rho_2} &= \frac{\beta}{3}\left(\rho_1 + \rho_2 + \rho_3 - 3\rho_0\right) + \frac{\beta}{3}\left(2\widetilde{\rho}_2 - \widetilde{\rho}_1 - \widetilde{\rho}_3\right) - \frac{\alpha_{\mathrm{v}}}{3}\left(\sigma_1 + \sigma_2 + \sigma_3\right) - \frac{\alpha_{\mathrm{d}}}{3}\left(2\widetilde{\sigma_2} - \widetilde{\sigma_1} - \widetilde{\sigma_3}\right) + \epsilon_2 \\ \frac{\partial\mathcal{U}_{\mathrm{cyt}}}{\partial\rho_3} &= \frac{\beta}{3}\left(\rho_1 + \rho_2 + \rho_3 - 3\rho_0\right) + \frac{\beta}{3}\left(2\widetilde{\rho}_3 - \widetilde{\rho}_1 - \widetilde{\rho}_2\right) - \frac{\alpha_{\mathrm{v}}}{3}\left(\sigma_1 + \sigma_2 + \sigma_3\right) - \frac{\alpha_{\mathrm{d}}}{3}\left(2\widetilde{\sigma_3} - \widetilde{\sigma_1} - \widetilde{\sigma_2}\right) + \epsilon_3 \\ \frac{\partial\mathcal{U}_{\mathrm{cyt}}}{\partial\epsilon_1} &= K\left(\epsilon_1 + \epsilon_2 + \epsilon_3\right) + \frac{2\mu}{3}\left(2\widetilde{\epsilon}_1 - \widetilde{\epsilon}_2 - \widetilde{\epsilon}_3\right) - \frac{1}{3}\left(\sigma_1 + \sigma_2 + \sigma_3\right) - \frac{1}{3}\left(2\widetilde{\sigma_1} - \widetilde{\sigma_2} - \widetilde{\sigma_3}\right) + \rho_1 \\ \frac{\partial\mathcal{U}_{\mathrm{cyt}}}{\partial\epsilon_2} &= K\left(\epsilon_1 + \epsilon_2 + \epsilon_3\right) + \frac{2\mu}{3}\left(2\widetilde{\epsilon}_2 - \widetilde{\epsilon}_1 - \widetilde{\epsilon}_3\right) - \frac{1}{3}\left(\sigma_1 + \sigma_2 + \sigma_3\right) - \frac{1}{3}\left(2\widetilde{\sigma_2} - \widetilde{\sigma_1} - \widetilde{\sigma_3}\right) + \rho_2 \\ \frac{\partial\mathcal{U}_{\mathrm{cyt}}}{\partial\epsilon_3} &= K\left(\epsilon_1 + \epsilon_2 + \epsilon_3\right) + \frac{2\mu}{3}\left(2\widetilde{\epsilon}_3 - \widetilde{\epsilon}_1 - \widetilde{\epsilon}_2\right) - \frac{1}{3}\left(\sigma_1 + \sigma_2 + \sigma_3\right) - \frac{1}{3}\left(2\widetilde{\sigma_3} - \widetilde{\sigma_1} - \widetilde{\sigma_2}\right) + \rho_3\end{aligned} \tag{S-13}$$

But

$$2\widetilde{x}_i - \widetilde{x}_j - \widetilde{x_k} = 3\widetilde{x}_i \tag{S-14}$$

with $(i, j, k)$ admitting right-hand permutation (scanning values 1,2,3). Therefore, at the steady state:

$$\begin{aligned}\frac{\partial\mathcal{U}_{\mathrm{cyt}}}{\partial\rho_i} &= \beta\left(\rho_i - \rho_0\right) - \left(\alpha_{\mathrm{v}}\sigma_{\mathrm{m}} + \alpha_{\mathrm{d}}\widetilde{\sigma}_i + \alpha_{\mathrm{a}}\delta_{i1}\sigma_i\right) + \epsilon_i = 0 \\ \frac{\partial\mathcal{U}_{\mathrm{cyt}}}{\partial\epsilon_i} &= \left(3K - 2\mu\right)\epsilon_{\mathrm{m}} + 2\mu\epsilon_i - \sigma_i + \rho_i = 0\end{aligned} \tag{S-15}$$

where $\sigma_\mathrm{m} = (\sigma_1 + \sigma_2 + \sigma_3)/3$ and $\epsilon_\mathrm{m} = (\epsilon_1 + \epsilon_2 + \epsilon_3)/3$ denote mean normal stress and strain, respectively.

Let's process the solution under the simple, yet prevalent, condition of $\alpha_\mathrm{v} = \alpha_\mathrm{d} = \alpha$. Therefore, Eq. (S-15)$_1$ would turn into

$$\frac{\partial \mathcal{U}_\mathrm{cyt}}{\partial \rho_i} = \beta(\rho_i - \rho_0) - \alpha_e \sigma_i + \epsilon_i = 0 \tag{S-16}$$

where $\alpha_e = \alpha + \alpha_\mathrm{a}\delta_{i1}$ is an effective anisotropic feedback parameter. Together with Eq. (S-15)$_2$, the following system of equations would emerge:

$$\begin{aligned} \rho_i - \frac{\alpha_e}{\beta}\sigma_i &= \rho_0 - \frac{\epsilon_i}{\beta} \\ -\rho_i + \sigma_i &= 3K\epsilon_\mathrm{m} + 2\mu\widetilde{\epsilon_i} \end{aligned} \tag{S-17}$$

which yields the following solution

$$\begin{aligned} \sigma_i &= 3\overline{K}^{(i)}\epsilon_\mathrm{m} + 2\overline{\mu}^{(i)}\widetilde{\epsilon_i} + \overline{\rho}_0^{(i)} \ \therefore\ \sigma_\mathrm{m} = 3\overline{K}_\mathrm{m}\epsilon_\mathrm{m} + \overline{\rho}_{0\mathrm{m}}\ ,\ \widetilde{\sigma_i} = 2\overline{\mu}^{(i)}\widetilde{\epsilon_i} \\ \rho_i &= 3\overline{K}_\rho^{(i)}\epsilon_\mathrm{m} + 2\overline{\mu}_\rho^{(i)}\widetilde{\epsilon_i} + \overline{\rho}_0^{(i)} \ \therefore\ \rho_\mathrm{m} = 3\overline{K}_{\rho\mathrm{m}}\epsilon_\mathrm{m} + \overline{\rho}_{0\mathrm{m}}\ ,\ \widetilde{\rho_i} = 2\overline{\mu}_\rho^{(i)}\widetilde{\epsilon_i} \end{aligned} \tag{S-18}$$

where $\bar{K}_\mathrm{m} = \left(\bar{K}^{(1)} + \bar{K}^{(2)} + \bar{K}^{(3)}\right)/3$, $\bar{K}_{\rho\mathrm{m}} = \left(\bar{K}_\rho^{(1)} + \bar{K}_\rho^{(2)} + \bar{K}_\rho^{(3)}\right)/3$, $\bar{\rho}_{0\mathrm{m}} = \left(\bar{\rho}_0^{(1)} + \bar{\rho}_0^{(2)} + \bar{\rho}_0^{(3)}\right)/3$ and the stiffness terms in general sense (with $\alpha_\mathrm{v} \neq \alpha_\mathrm{d}$) read

$$\begin{aligned} &3\bar{K}^{(i)} = \frac{3K\beta - 1}{\beta - \alpha_\mathrm{ev}^{(i)}} \quad , \quad 3\bar{K}_\rho^{(i)} = \frac{3K\alpha_\mathrm{ev}^{(i)} - 1}{\beta - \alpha_\mathrm{ev}^{(i)}} \quad , \quad \alpha_\mathrm{ev}^{(i)} = \alpha_\mathrm{v} + \alpha_\mathrm{a}\delta_{i1} \\ &2\bar{\mu}^{(i)} = \frac{2\mu\beta - 1}{\beta - \alpha_\mathrm{ed}^{(i)}} \quad , \quad 2\bar{\mu}_\rho^{(i)} = \frac{2\mu\alpha_\mathrm{ed}^{(i)} - 1}{\beta - \alpha_\mathrm{ed}^{(i)}} \quad , \quad \alpha_\mathrm{ed}^{(i)} = \alpha_\mathrm{d} + \alpha_\mathrm{a}\delta_{i1} \\ &\bar{\rho}_0^{(i)} = \frac{\beta}{\beta - \alpha_\mathrm{ev}^{(i)}}\rho_0 \end{aligned} \tag{S-19}$$

##### 1.2 Nonlinear anisotropic model

Recalling Eq. (S-4), the cell active free energy is superposed by a nonconservative component from ATP consumption:

$$\mathcal{U}_\mathrm{cell} = \mathcal{U}_\mathrm{act} + \mathcal{U}_\mathrm{pas} - \int_{\xi_i=0}^{\rho_i} \kappa(\sigma_i)\,\Delta G \mathrm{d}\xi_i \tag{S-20}$$

where

$$\kappa(\sigma) = \frac{(\alpha\sigma)^p}{1 + c(\alpha\sigma)^p}$$

Noting that the nonconservative term in Eq. (S-20) replaces the chemomechanical energy portion of $\mathcal{U}_\mathrm{chem}$ in Eq. (S-10), the total cytoskeletal energy density can be expressed in the following expanded form:

$$\begin{aligned} \mathcal{U}_\mathrm{cyt} = &\frac{K}{2}(\epsilon_1 + \epsilon_2 + \epsilon_3)^2 + \mu\left(\widetilde{\epsilon_1}^2 + \widetilde{\epsilon_2}^2 + \widetilde{\epsilon_3}^2\right) - \frac{1}{6}(\sigma_1 + \sigma_2 + \sigma_3)(\epsilon_1 + \epsilon_2 + \epsilon_3) \\ &- \frac{1}{2}\left(\widetilde{\sigma_1}\widetilde{\epsilon_1} + \widetilde{\sigma_2}\widetilde{\epsilon_2} + \widetilde{\sigma_3}\widetilde{\epsilon_3}\right) + \frac{\beta}{6}(\rho_1 + \rho_2 + \rho_3 - 3\rho_0)^2 + \frac{\beta}{2}\left(\widetilde{\rho_1}^2 + \widetilde{\rho_2}^2 + \widetilde{\rho_3}^2\right) + \\ &\rho_1\epsilon_1 + \rho_2\epsilon_2 + \rho_3\epsilon_3 - \frac{1}{2}\Delta G\left[\kappa(\sigma_1)\rho_1 + \kappa(\sigma_2)\rho_2 + \kappa(\sigma_3)\rho_3\right] \end{aligned} \tag{S-21}$$

Using the same identity as reflected in Eq. (S-12), one can write

$$\begin{aligned}
\frac{\partial \mathcal{U}_{\mathrm{cyt}}}{\partial \rho_1} &= \frac{\beta}{3}\left(\rho_1+\rho_2+\rho_3-3\rho_0\right)+\frac{\beta}{3}\left(2\widetilde{\rho_1}-\widetilde{\rho_2}-\widetilde{\rho_3}\right)+\epsilon_1-\kappa\left(\sigma_1\right)\Delta G \\
\frac{\partial \mathcal{U}_{\mathrm{cyt}}}{\partial \rho_2} &= \frac{\beta}{3}\left(\rho_1+\rho_2+\rho_3-3\rho_0\right)+\frac{\beta}{3}\left(2\widetilde{\rho_2}-\widetilde{\rho_1}-\widetilde{\rho_3}\right)+\epsilon_2-\kappa\left(\sigma_2\right)\Delta G \\
\frac{\partial \mathcal{U}_{\mathrm{cyt}}}{\partial \rho_3} &= \frac{\beta}{3}\left(\rho_1+\rho_2+\rho_3-3\rho_0\right)+\frac{\beta}{3}\left(2\widetilde{\rho_3}-\widetilde{\rho_1}-\widetilde{\rho_2}\right)+\epsilon_3-\kappa\left(\sigma_3\right)\Delta G \\
\frac{\partial \mathcal{U}_{\mathrm{cyt}}}{\partial \epsilon_1} &= K\left(\epsilon_1+\epsilon_2+\epsilon_3\right)+\frac{2\mu}{3}\left(2\widetilde{\epsilon_1}-\widetilde{\epsilon_2}-\widetilde{\epsilon_3}\right)-\frac{1}{3}\left(\sigma_1+\sigma_2+\sigma_3\right)-\frac{1}{3}\left(2\widetilde{\sigma_1}-\widetilde{\sigma_2}-\widetilde{\sigma_3}\right)+\rho_1 \\
\frac{\partial \mathcal{U}_{\mathrm{cyt}}}{\partial \epsilon_2} &= K\left(\epsilon_1+\epsilon_2+\epsilon_3\right)+\frac{2\mu}{3}\left(2\widetilde{\epsilon_2}-\widetilde{\epsilon_1}-\widetilde{\epsilon_3}\right)-\frac{1}{3}\left(\sigma_1+\sigma_2+\sigma_3\right)-\frac{1}{3}\left(2\widetilde{\sigma_2}-\widetilde{\sigma_1}-\widetilde{\sigma_3}\right)+\rho_2 \\
\frac{\partial \mathcal{U}_{\mathrm{cyt}}}{\partial \epsilon_3} &= K\left(\epsilon_1+\epsilon_2+\epsilon_3\right)+\frac{2\mu}{3}\left(2\widetilde{\epsilon_3}-\widetilde{\epsilon_1}-\widetilde{\epsilon_2}\right)-\frac{1}{3}\left(\sigma_1+\sigma_2+\sigma_3\right)-\frac{1}{3}\left(2\widetilde{\sigma_3}-\widetilde{\sigma_1}-\widetilde{\sigma_2}\right)+\rho_3
\end{aligned} \tag{S-22}$$

which, upon secondary reference to Eq. (S-14), takes the following encapsulated form at the steady state:

$$\begin{aligned}
\frac{\partial \mathcal{U}_{\mathrm{cyt}}}{\partial \rho_i} &= \beta\left(\rho_i-\rho_0\right)-\kappa(\sigma_i)\Delta G+\epsilon_i=0 \\
\frac{\partial \mathcal{U}_{\mathrm{cyt}}}{\partial \epsilon_i} &= \left(3K-2\mu\right)\epsilon_{\mathrm{m}}+2\mu\epsilon_i-\sigma_i+\rho_i=0
\end{aligned} \tag{S-23}$$

where $\sigma_{\mathrm{m}}=\left(\sigma_1+\sigma_2+\sigma_3\right)/3$ and $\epsilon_{\mathrm{m}}=\left(\epsilon_1+\epsilon_2+\epsilon_3\right)/3$.

Equation (S-23) can be further simplified into the following nonlinear system of equations:

$$\begin{cases}
\sigma_i-\frac{\kappa(\sigma_i)}{\beta'}=\rho_0+3K'\epsilon_{\mathrm{p}}+2\mu'\widetilde{\epsilon_i} \\
\rho_i-\frac{\kappa(\sigma_i)}{\beta'}=\rho_0-\left(3K'_\rho\epsilon_{\mathrm{p}}+2\mu'_\rho\widetilde{\epsilon_i}\right)
\end{cases} \tag{S-24}$$

which replicates Eq. [4] of the manuscript for ease of reference. For easier reference, $3K'=3K-1/\beta$, $2\mu'=2\mu-1/\beta$ and $3K'_\rho=2\mu'_\rho=1/\beta$ define the effective bulk and shear moduli and their counterparts for motor density, respectively. Moreover, $\beta'=\beta/\Delta G$, and $\kappa$ was defined in Eq. (S-20) as function of principal stresses.

Having obtained their principal components, the corresponding tensorial representation of stress and contractility can be determined from the polar decomposition theory as

$$\begin{cases}
\boldsymbol{\sigma}=\sigma_i\boldsymbol{n}^{(\mathrm{i})}\otimes\boldsymbol{n}^{(\mathrm{i})} \\
\boldsymbol{\rho}=\rho_i\boldsymbol{n}^{(\mathrm{i})}\otimes\boldsymbol{n}^{(\mathrm{i})}
\end{cases} \tag{S-25}$$

where $\boldsymbol{n}^{(\mathrm{i})}$ denote the principal directions, and summation is implied over $i$.

#### 2 Model Predictions

The total free energy is superposed from three sources: cell, cell-ECM interface, and ECM itself, hence reading $\mathcal{U}_{\mathrm{tot}}=\mathcal{U}_{\mathrm{cell}}+\mathcal{U}_{\mathrm{int}}+\mathcal{U}_{\mathrm{M}}$. $\mathcal{U}_{\mathrm{cell}}$ was formulated in Eq's (S-6) and (S-20) for the linear and nonlinear anisotropic models, respectively. While the cytoskeletal portions at steady state were expended in Eq's (S-11) and (S-21), the cell is presumably composed of the nucleus (idealized with a central sphere), nesprin (idelized with an interfacial layer between the nucleus and cytoskeleton) and cytoskeleton (schematized in

Fig. 2 of the manuscript). Hence, the passive energy must comprise the strain energy stored in all the above when deformed by contractile forces. That is

$$\mathcal{U}_{\text{pas}} = \int_{\Omega_{\text{nuc}}} W_{\text{nuc}}\left(\epsilon_{ij}\right) \mathrm{d}\Omega + \int_{\Omega_{\text{nes}}} W_{\text{nes}}\left(\epsilon_{ij}\right) \mathrm{d}\Omega + \int_{\Omega_{\text{cyt}}} W_{\text{cyt}}\left(\epsilon_{ij}\right) \mathrm{d}\Omega \tag{S-26}$$

where $W$'s denote the respective strain energy densities as functions of strain.

Moreover, interfacial energy represents the contributions from adhesions (mainly integrins), actin cortex and the basement membrane. The interface is here idealized as a thin layer with uniform thickness that connects the cytoskeleton to the matrix (see Fig. 1a in the manuscript). Adhesion receptors (mainly integrins) maintain spontaneous binding to complimentary collagen ligands, which results in reduced free energy [13]. Hence, the cell tends to adopt an elongated shape at higher receptor densities to maximize the number of bonds with the matrix. On the other hand, the membrane-cortical tension together tend to shrink the surface area, not favoring cell elongation. Altogether, the combined effect of membrane-cortical tension and adhesion receptors can be expressed in terms of a factor of cell body surface area as

$$U_{\text{int}} = \left(\gamma_0 - \gamma_1 \delta_{\text{ad}}\right) \mathcal{S} \tag{S-27}$$

with $\gamma_0$ denoting the contribution of membrane-cortical tension, $\gamma_1$ reflecting the change of energy associated with the formation of each receptor-ligand bond, and $\delta_{\text{ad}}$ being a scaling factor representing the density of adhesion receptors. Finally, $\mathcal{S}$ represents the cell outer surface area which, for a prolate spheroidal cell, obeys

$$\mathcal{S} = 2\pi b^2 \left(1 + \frac{\zeta}{e} \arcsin e\right)$$

where $\zeta = a/b \geqslant 1$ is the cell aspect ratio (with $\zeta \geqslant 1$ being characteristic of a prolate cell) and $e^2 = 1 - \frac{1}{\zeta^2}$ denotes the spheroid eccentricity.

Finally, matrix energy is constituted partially by the strain energy stored in ECM by the cell's contractile forces (through adhesion receptors) and partially by fibers. The latter is contributed by diffuse and aligned fibers. Diffuse fibers are randomly dispersed throughout the matrix and buckled along the principal compressive stresses while aligned fibers are stretched along the tensile principal stresses. Accordingly, the combined energy contributed by ECM fibers can be considered to admit a nearly incompressible Neo-Hookean constitutive model with a quadratic-type energy expression [14]. That is, the combined ECM strain energy density reads

$$W_{\text{M}} = \frac{\mu_{\text{M}}}{2}\left(\frac{\sum_{i=1}^{3}\lambda_i^2}{\left(\lambda_1\lambda_2\lambda_3\right)^{2/3}} - 3\right) + \frac{K_{\text{M}}}{2}\left(\lambda_1\lambda_2\lambda_3 - 1\right)^2 + \sum_{i=1}^{3} f\left(\lambda_i\right) \tag{S-28}$$

where the last term is provided by aligned fibers and the previous terms represent randomly dispersed fibers. Here, $\mu_{\text{M}}$ and $K_{\text{M}}$ denote the matrix shear and bulk moduli, and $\lambda_i$ $(i = 1, 2, 3)$ are the three principal stretches. The power-law function $f$, hereby introduced to capture fiber-mediated matrix stiffening in the tensile principal directions, is a piecewise function obeying

$$f\left(\lambda_i\right) = \begin{cases} 0 \quad , \quad \lambda_i < \lambda_{\text{L}} \\ E_{\text{f}} \frac{(\lambda_i - \lambda_{\text{L}})^{n+2}(\lambda_{\text{U}} - \lambda_{\text{L}})^{-n}}{(n+1)(n+2)}, \lambda_{\text{L}} < \lambda_i < \lambda_{\text{U}} \\ E_{\text{f}}\left[\frac{\lambda_{\text{U}}(\lambda_i - \frac{1}{2}\lambda_i^2)}{n+1} + \frac{(1-\lambda_{\text{U}}+\lambda_i)^{m+2} - (m+2)\lambda_i}{(m+1)(m+2)}\right], \lambda_i > \lambda_{\text{U}} \end{cases} \tag{S-29}$$

where $E_{\mathrm{f}}$ denotes fibers' Young's modulus, and

$$(\lambda_{\mathrm{L}}, \lambda_{\mathrm{U}}) = \lambda_{\mathrm{c}} \mp \frac{\lambda_{\mathrm{c}} - 1}{8}$$

where $\lambda_{\mathrm{L}}$ (using $-$) and $\lambda_{\mathrm{U}}$ (using $+$) are, respectively, the lower and upper bounds to the interpolation range, so formulated as to ensure a continuous derivative around the transition point identified by the critical stretch $\lambda_{\mathrm{c}}$. Correspondingly, $f$ is primarily aimed to distinguish between loose (randomly oriented) fibers (with vanishing principal stresses) from aligned fibers (with power-like hardening), with the transition in between determined by $\lambda_{\mathrm{c}}$.

Prediction of the steady-state optimum cell body aspect ratio solidifies into the competition between decreasing cell-ECM and increasing interfacial energy components, leading to a non-monotonic profile of the total free energy $\Delta\mathcal{U}_{\mathrm{tot}}$ (here presented in terms of its difference with the value corresponding to a spherical cell, *i.e.* $\Delta\mathcal{U} \equiv \mathcal{U} - \mathcal{U}(\zeta = 1)$) with respect to the cell body aspect ratio $\zeta$. Evaluation of $\zeta_{\mathrm{opt}}$ then results from the postprocessing of the 3D implementation of the model, where the nonlinear constitutive model was imposed in COMSOL 5.6 so as to be solved for stress and contractility components at every time step. Each point on the $\zeta_{\mathrm{opt}}$ plots would then stem from simultaneous solution of equilibrium and constitutive equations scanned throughout a prescribed range of cell body aspect ratios $1 \leqslant \zeta \leqslant 20$. Energy differences $\Delta U_{\mathrm{tot}}$ were then exported to a post-processing subroutine to evaluate the optimum aspect ratio $\zeta_{\mathrm{opt}}$ from the local minimum to the $\Delta U_{\mathrm{tot}}$ vs. $\zeta$ curve, as showcased in Fig's S-1 and S-4.

#### 2.1 Linear anisotropic model

Figure S-1 reveals the volumetric integral of the total cell energy with stress and contractility obeying the linear model as culminated in Eq. (S-24). Accordingly, the optimum cell aspect ratios collected at varying steady states as function of the ECM-to-cell stiffness ratio ($\eta = E_{\mathrm{M}}/E_{\mathrm{C}}$) are shown in Fig's S-1 (b) and (c) in terms of the normalized adhesion density and anisotropic chemomechanical coupling parameter $\alpha_{\mathrm{a}}$, respectively.

As revealed by Fig. S-1, the linear anisotropic model truly suggests a spherical shape domination in both extremely compliant and stiff matrices. Nevertheless, the shape transition from soft to stiff matrices is abrupt (*i.e.* of the first order). Recent experimental studies, however, transpire the existence of a transition zone with a highly heterogeneous distribution dominated by both spherical and elongated cell shapes [15]. Such lacking capability within the linear model warrants considering a trizonal model which can capture the overly soft and stiff as well as the meso-stiff matrix zone as presented in the manuscript.

#### 2.2 Nonlinear anisotropic model

##### 2.2.1 One-dimensional special case

Recollecting Eq. (S-24)$_1$, only one independent or nonzero stress-strain conjugate exists in the 1D special case. That is, assuming the nonzero stress-strain conjugate at the cell-post interface or the uniform stress-strain conjugate at the cell-ECM interface is denoted with $\sigma$ and $\epsilon$, the correlation obeys $\epsilon = -2\sigma/K_{\mathrm{p}}$ and $\epsilon \equiv \epsilon_{\mathrm{rr}}(R) = \epsilon_{\theta\theta}(R) = -\sigma/2\mu_{\mathrm{M}}$, respectively. For the latter case, the reader can consult the solution for a sphere with internal pressure [16]. Therefore, the general 1D model solidifies into the solution to the equation identified by $Y_1 = Y_2$, where

$$\begin{cases} Y_1 = \dfrac{\kappa\left(\frac{\sigma}{\rho_0}\right)}{\rho_0\beta'} \\ \\ Y_2 = -1 + \left(1 + \frac{\gamma\mathcal{G}}{\eta}\right)\frac{\sigma}{\rho_0} \end{cases} \tag{S-30}$$

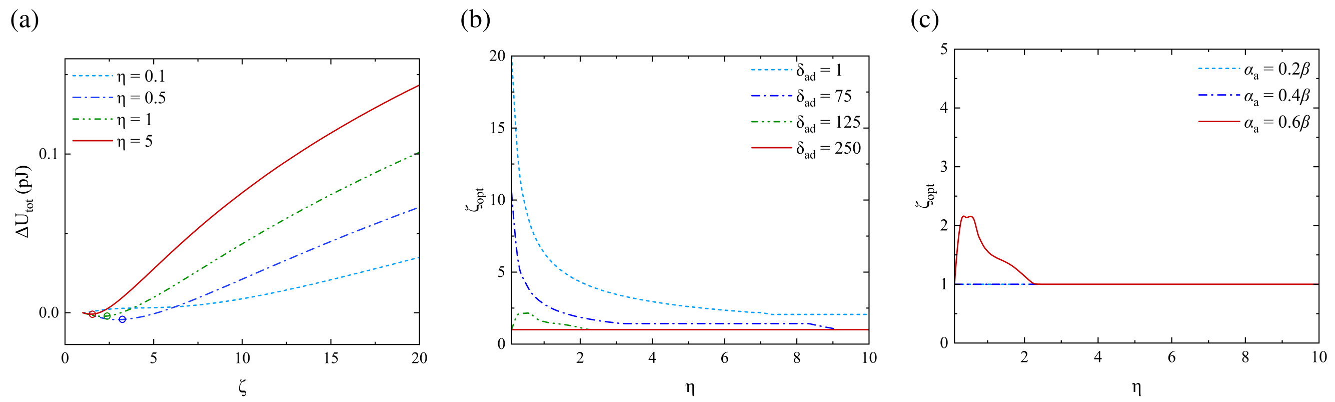


Figure S-1: **Linear anisotropic optimum cell body aspect ratio vs. ECM stiffness.** Variation of optimum cell body aspect ratio as function of the ECM to cell stiffness ratio $\eta \equiv E_{\mathrm{M}}/E$ predicted from the linear anisotropic constitutive model in Eq. (S-24): (a) total cell free energy (in difference with that of the spherical cell) vs. cell body aspect ratio $\zeta$ for various ECM to cell stiffness ratios; (b) optimum cell body aspect ratio vs. $\eta$ at varying integrin contribution $\delta_{\mathrm{ad}}$ (at fixed membrane–cortical tensile stiffness $\gamma_0 = 0.2$ mN/m); (c) at varying anisotropic chemomechanical coupling parameter $\alpha_{\mathrm{a}}$ (at fixed values of $\gamma_0 = 0.2$ mN/m and $\delta_{\mathrm{ad}} = 1$). Other cell parameters are identified by $E = 1.5$ kPa, $\nu = 0.27$ and $(\varsigma \equiv \rho_0/E, \mathcal{B} \equiv \beta E) = (0.2, 4.5)$. Adhesion binding affinity stiffness is also fixed at $\gamma_1 = 0.001$ mN/m.

where $\gamma$ and $\mathcal{G} = g - 1/\mathcal{B}$ are problem-dependent dimensionless parameters, such that

$$\gamma = 2 \quad , \quad g = \frac{1-\nu}{(1+\nu)(1-2\nu)}$$

for the cell connected to microposts, and

$$\gamma = 1+\nu \quad , \quad g = 1-2\nu$$

for the spherical cell connected to infinite ECM. Moreover, $\mathcal{B} = \beta E$ and the rest of parameters were defined in advance.

In the general sense, Eq. (S-30) delivers a spectral solution comprising one, two or three solution branches. Accordingly, the solution branch is governed by casewise comparison between the line slope $(1/\eta)$ with the two slopes of tangency between $Y_1$ and $Y_2$, here denoted with critical slopes $\eta_{\mathrm{c}}^{(1)}$ and $\eta_{\mathrm{c}}^{(2)}$ as schematized in Fig. S-2c. At the points of tangency, simultaneous satisfaction of $Y_1\left(\frac{\sigma}{\rho_0}\right) = Y_2\left(\frac{\sigma}{\rho_0}\right)$ and ${Y'}_1\left(\frac{\sigma}{\rho_0}\right) = {Y'}_2\left(\frac{\sigma}{\rho_0}\right)$, utilizing $\frac{\sigma}{\rho_0} = \frac{1}{\alpha\rho_0}\left(\frac{\kappa}{1-c\kappa}\right)^{1/p}$ which expresses $\sigma$ in terms of $\kappa$, the following equation can be reached:

$$\begin{aligned}
&\kappa_{\mathrm{c}}^2 - q'\kappa_{\mathrm{c}} + {\rho\prime}_0\beta'(1-q) = 0 \ \therefore \ \kappa_{\mathrm{c}} = \frac{1}{2}\left(q' \pm \sqrt{\Delta}\right) = \{\kappa_{\mathrm{c}}^{(1)}, \kappa_{\mathrm{c}}^{(2)}\} \\
&\eta_{\mathrm{c}} = \frac{\gamma\mathcal{G}}{-1 + \frac{p\alpha}{\beta'}{\kappa_{\mathrm{c}}}^q(1-c\kappa_{\mathrm{c}})^{2-q}} = \{\eta_{\mathrm{c}}^{(1)}, \eta_{\mathrm{c}}^{(2)}\}
\end{aligned} \tag{S-31}$$

where $\Delta = {q'}^2 - \frac{4}{p}{\rho\prime}_0\beta'$ is the equation discriminant, with $q = 1 - 1/p$ and $\left(q', {\rho'}_0\right) = \frac{1}{c}(q, \rho_0)$.

Figure S-2 shows a schematic representation of the 1D solution through varying intersection between $Y_1$ and $Y_2$, mathematical notion of the critical stiffness ratios corresponding to tangential contact between

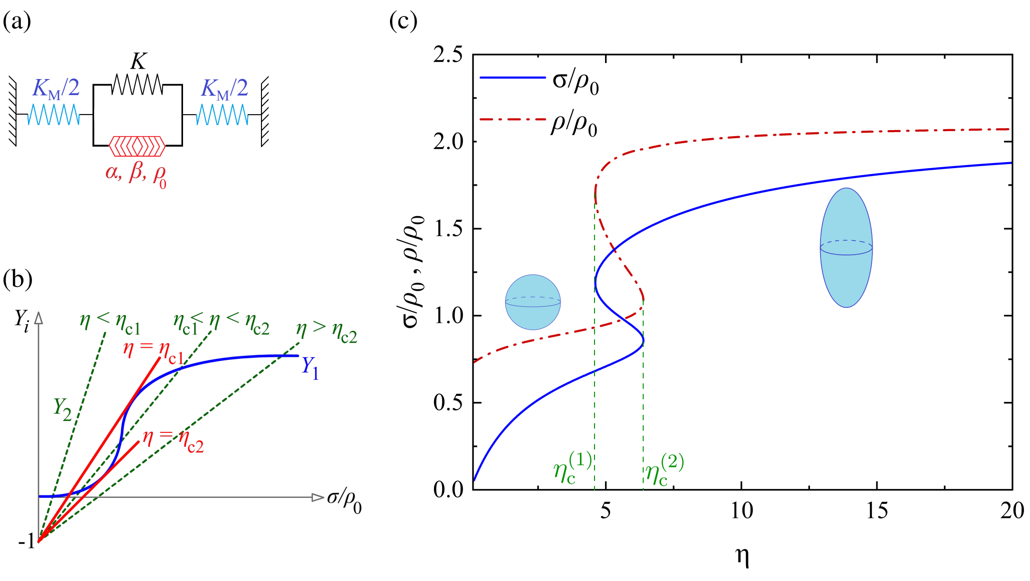


Figure S-2: **Mathematical representation of the trizonal chemomechanical model.** (a) 1D cell-ECM model consisting of linear elements (passive cell stiffness attached to ECM stiffness) and a nonlinear active element; (b) schematic 1D solution scheme from intersection between the stress-dependant nonlinear term and stiffness-dependant linear term, revealing two critical stiffness ratios $\eta_c^{(1)}$ and $\eta_c^{(2)}$ determining the bounds to solution branches; (c) example complete 1D solution (with $p = 8$) delivering stress and contractility over three branches, with cell and parameters identified by $E = 2$ kPa, $\nu = 0.3$, $(\varsigma \equiv \rho_0/E, \mathcal{B} \equiv \beta E, \Lambda \equiv \alpha/\beta) = (0.15, 6, 1)$, which delivers $\rho_0 = 0.3$ kPa $\beta = 3$ 1/kPa and $\alpha = \beta$.

$Y_1$ and $Y_2$, and an example trizonal solution for stress and contractility vs. stiffness ratio $\eta$. The existence of two distinct motility levels, with the upper level exceeding twice the lower level with the adopted parameters, is conspicuous. Upon generalization of the model into 3D domains, the lower and upper branches are predominantly conducive to spherical (non-invasive) and elongated (metastatic) cells, respectively. Note that, albeit mathematically true, the middle declining portion in the overlapping (middle) branch is nonphysical considering the essential condition of stress and contractility elevation with the matrix being stiffened. Hence, the middle solution will be disregarded in the general (3D) model implementation.

Figure S-3 shows example plots of trizonal solution for stress and contractility vs. stiffness ratio $\eta$ at varying modeling tuning parameters $p$ and $c$ for the special case of a cell connected to microposts. In order for better consistency with the general case, the post stiffness is conceived in terms of an equivalent matrix bulk modulus, whose equivalent Young's modulus is identified in proportion to that of the cell as $E_{\mathrm{M}} = \eta E$, hence letting $K_{\mathrm{p}} = E_{\mathrm{M}}/3(1 - 2\mu_{\mathrm{M}})$. Note anew that the linear model is retrievable by letting $(p, c) = (1, 0)$. Increasing exponent $p$ would chiefly decrease the upper-branch motility and contractile force level while increasing the unbinding-to-binding rate ratio $c$ would would shrink the stiffness overlapping regime conducive to two possible cell shapes. Hence, equal binding and unbinding rates ($c = 1$) yields a neutral state with no or little distinction between lower and upper-branch motility levels and cell shapes. Altogether, tuning parameters best commensurate with physics of cell shape transition should lie within the ranges of $4 \leqslant p \leqslant 10$ and $0.3 \leqslant c < 0.6$.

##### *2.2.2 Application to 3D ellipsoidal cells: Energetics-based optimum cell shape*

Figure S-4 showcases variations in the energy profile predicted from the lower and upper branches of the nonlinear model (see Fig. 3 of the manuscript for clarity) under the effect of matrix stiffening (a,b) as well

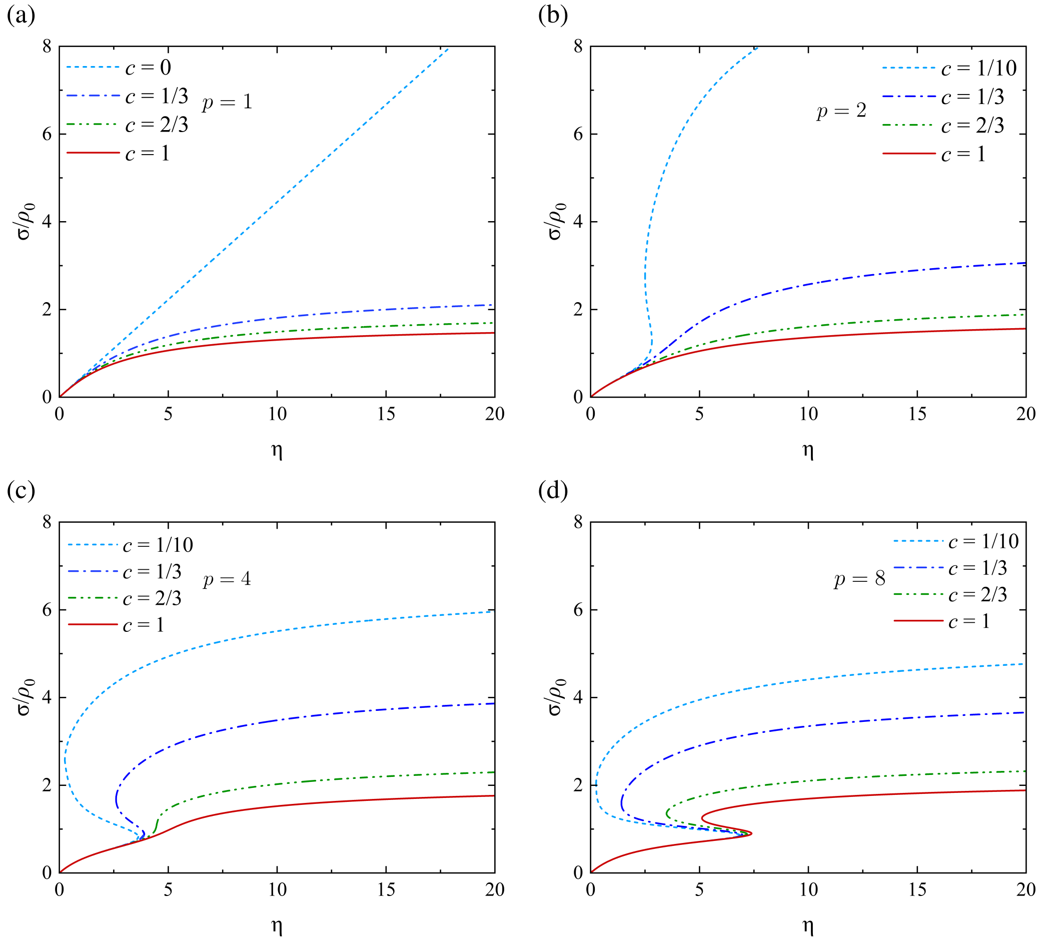


Figure S-3: **Trizonal solution at varying tuning parameters** $(p, c)$**.** One-dimensional trizonal solution for normalized stress vs. equivalent stiffness ratio $\eta \equiv E_{\mathrm{p}}/E$ at various tuning parameters of $p$ ranging from 1 to 8 and $c$ adopting (nearly) zero to 1. Cell parameters are identified by $E = 1.5$ kPa, $\nu = 0.27$, $(\varsigma, \mathcal{B}, \Lambda) = (0.2, 4.5, 1)$.

as membrane-cortical stiffening (c,d) and integrin intensification (e,f). The increasing effect of stiffness on the ECM strain energy would transpire as maintained optimum aspect ratio of 1 in the lower branch and reducing optimum aspect ratio towards 1 in the upper branch.

The energy trends predicted by Fig. S-4 curves are fully commensurate with physics of cell–ECM interaction with increasing elongation/decreasing circularity. First and foremost, the passive part and hence the total free energy is intuitively elevated with increasing matrix stiffness. Note that, due to its limited ATP consumption and contractility, the lower branch of the constitutive model can only predict (nearly) spherical cell shapes. Hence, $\mathcal{U}_{\text{tot}}$ corresponding to the lower branch is normally a monotonically increasing function of cell body aspect ratio, thus its optimum realized at $\zeta = 1$. While the lower branch admits increasing level of total energy and ATP consumption with increasing ECM and cell body aspect ratio, the upper branch admits craters within the energy contours representing optimal cell body aspect ratio decreasing with stiffness. This extended trend is well revealed by plots of the optimum aspect ratio against the ECM normalized stiffness in Fig. 3 of the manuscript.

Next, varying membrane-cortical tension and adhesion density is sensibly observable in Fig's S-4 (c,d) and (e,f), respectively. Intuitively, increasing the former would further withstand cell elongation due to higher membrane stiffness. All the same, increasing the latter would impair the cell's ability of adherance

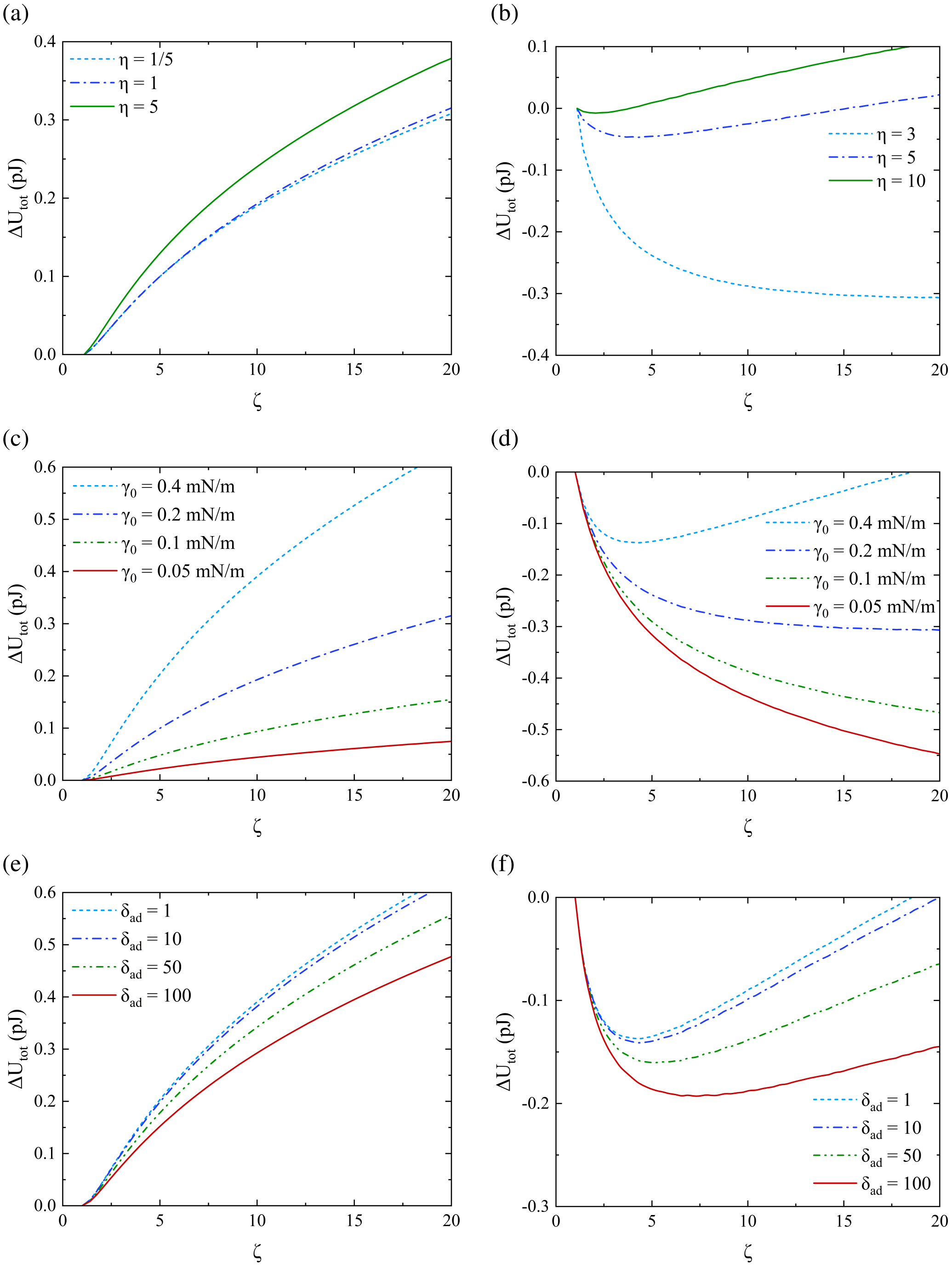


Figure S-4: **Cell–ECM total free energy predicted by the nonlinear model as function of cell shape.** (a,b) Variation of cell–ECM free energy difference predicted by the lower and upper branches vs. cell body aspect ratio $\zeta$ at select ECM to cell stiffness ratios $\eta \equiv E_{\mathrm{M}}/E$; (c,d) similar variation at select membrane-cortical stiffnesses $\gamma_0$ (at fixed integrin contribution $\delta_{\mathrm{ad}} = 1$); (e,f) similar variation at select integrin contribution $\delta_{\mathrm{ad}}$ (at fixed membrane-cortical stiffnesses $\gamma_0 = 0.2$ mN/m). Cell parameters are identified by $E = 1.5$ kPa, $\nu = 0.27$, $(\varsigma, \mathcal{B}, \Lambda) = (0.2, 5, 0.7)$. Also unless stated otherwise, $(\gamma_0, \gamma_1) = (0.2, 0.001)$ mN/m and $\delta_{\mathrm{ad}} = 1$. Model tuning parameters are also fixed at $(p, c) = (6, 1/3)$.

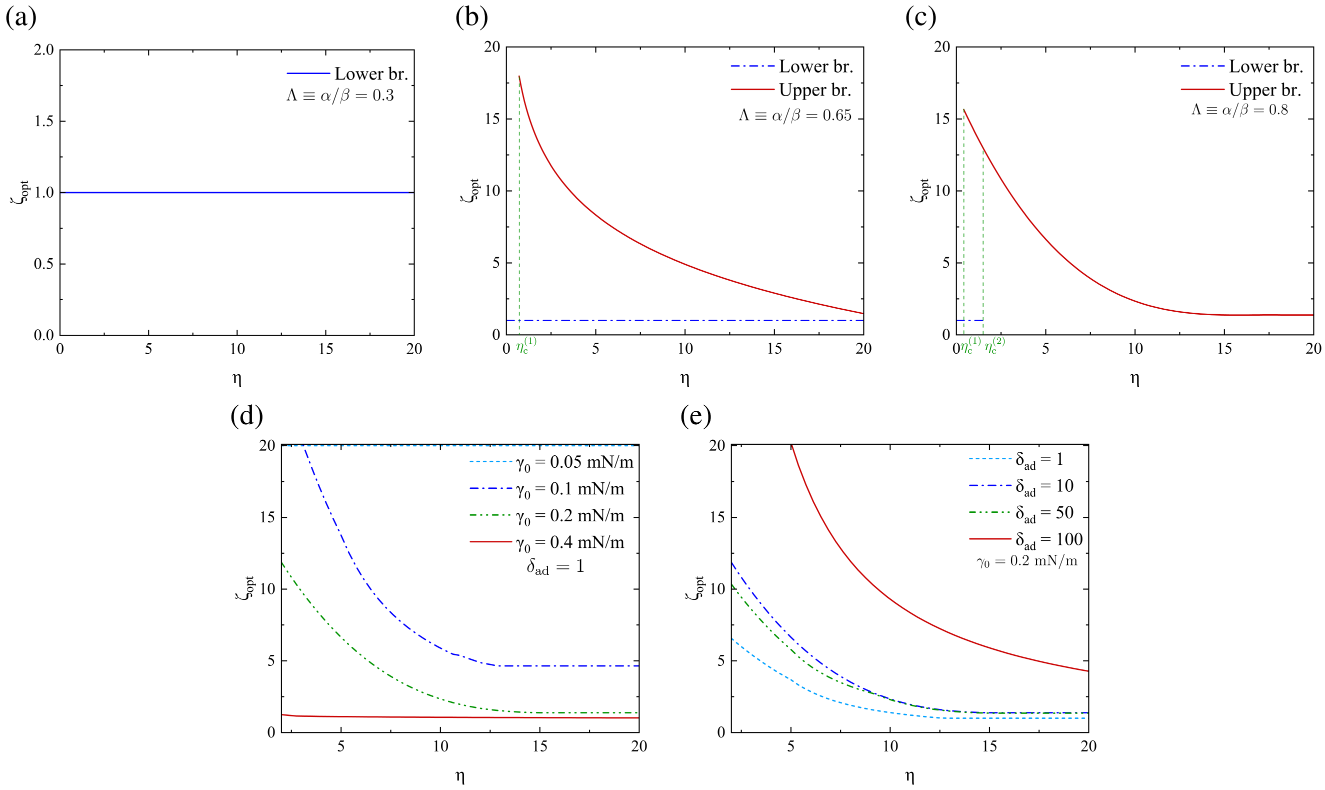


Figure S-5: **Optimum cell body aspect ratio vs. ECM stiffness.** Variation of lower vs. upper-branch optimum cell body aspect ratio as function of the ECM to cell stiffness ratio $\eta \equiv E_{\mathrm{M}}/E$: (a–c) complete solution at various normalized polarization parameters $\Lambda \equiv \alpha/\beta$; (d,e) upper-branch (highly motile) solution at varying membrane–cortical tensile stiffness $\gamma_0$ (with $\delta_{\mathrm{ad}} = 1$) and integrin contribution $\delta_{\mathrm{ad}}$ (with $\gamma_0 = 0.2$ mN/m), respectively. Other cell parameters are identified by $E = 1.5$ kPa, $\nu = 0.27$ and $(\varsigma, \mathcal{B}) = (0.2, 4.5)$. Adhesion binding affinity stiffness is fixed at $\gamma_1 = 0.001$ mN/m. Model tuning parameters are also fixed at $(p, c) = (6, 1/3)$.

to and interaction with the matrix, hence downregulating actomyosin recruitment and inhibiting ATP consumption [17, 18, 19, 20, 21].

The predicted optimum aspect ratios evaluated at the local minima observable in Fig. S-4 are next collected in plots against the ECM normalized stiffness $\eta$ in Fig. S-5. The normalized polarization parameter $\Lambda \equiv \alpha/\beta$, which quantifies the intensity of stress-regulated signaling pathways, would render the optimum steady-state cell body aspect ratio more sensitive to ECM stiffening. Moreover, it would reduce heterogeneity in cell motility by shrinking the overlapping range of $\eta$ admitting heterogeneous shape distribution. Namely, larger stress polarization would clearly lower the minimum critical ECM stiffness that triggers metastasis. Smaller than infimum polarization parameters, however, would represent signaling detachment between the cell and ECM (Fig. S-5a), hence predicting non-invasive spherical cells regardless of ECM stiffness.

Moreover, the same variation at varying membrane-cortical tension and adhesion density can be seen in Fig. S-5d,S-5e. Intuitively, increasing the former would further withstand cell elongation due to higher membrane stiffness. Vice versa, increasing the latter would impair the cell's ability of adherance to and interaction with the matrix, hence downregulating actomyosin recruitment and inhibiting ATP consumption [17, 18, 19, 20, 21]. Figure S-6 sheds further light on the effects of model parameters on the optimal cell shape. Notably, the decreasing trend within $\zeta_{\text{opt}}$ vs. $\eta$ might appear as contradictory with studies suggestive of increasing cell elongation and invasiveness with increasing ECM stiffness [22,23,24,9,25]. While all such studies are enlightening within their own confines, they all work around the transition zone, *i.e.* from below $\eta_c^{(1)}$ up to slightly above this critical limit. Rather, only studies unfolding the biphasic correlation between cell invasiveness and ECM stiffness have posed the full-range correlation, yet failed to predict it analytically [13, 26, 27]. The apparent controversy between the two observed trends cannot be duly reconciled unless by taking into account the effect of heterogeneity in actomyosin recruitment at fixed motor density. Such heterogeneity has been unveiled in some relevant studies, yet without any effort for its analytical prediction [28, 29, 30, 15].

Since the lower-branch solution prevalently delivers a spherical cell, the effect of model parameters can be explored for a sufficiently stiff matrix taken into account, *e.g.* for $\eta = 5$. Figure S-6 presents the upper-branch steady-state optimum cell body aspect ratio $\zeta_{\text{opt}}$ in terms of normalized model parameters quantifying stress polarization ($\Lambda \equiv \alpha/\beta$), contractile density ($\varsigma \equiv \rho_0/E$) and chemo-mechanical coupling ($\mathcal{B} \equiv \beta E$) as well as interfacial parameters including membrane-cortical tensile stiffness $\gamma_0$ (mN/m) and normalized adhesion (integrin) density $\delta_{\text{ad}}$.

As discussed beneath Fig's S-5(a–c), polarization parameters below the infimum threshold represent signaling detachment between the cell and ECM while higher-than-infimum values are indicative of active ATP consumption. Nevertheless, larger values of $\Lambda$ would trigger a steady state tending towards a stationary condition with small or zero myosin unbinding and net ATP consumption. This can be better perceived by referring to the primitive definition of $\alpha$, which sets a nonlinear correlation between binding and unbinding myosin motor rates, *i.e.* $k_{\text{on}} = (\alpha\sigma)^p\, k_{\text{off}}$. As such, extreme enlargement of $\alpha$ would prohibitively enlarge the myosin binding rate so that ATP consumption and mechanosensitive signaling between the cell and ECM would be inhibited. Further, the antithetical influences of the interfacial energy terms can be envisaged in Fig. S-6c,S-6d. It is witnessed that, below a certain limit, the tensile softening of the membrane would associate with significant increase in cell elongation and invasiveness. This condition represents membrane damage triggered by effects such as oxidative radiation [31], phenolic acids [32], or metal release processes [33]. All the same, the effect induced by integrin density is observed to escalate remarkably beyond a certain upper threshold that decreases considerably with membrane softening, *i.e.* lowering $\gamma_0$. Such abrupt increase in phenotype activity by increasing integrin contribution beyond this threshold is corroborated by earlier studies [17, 18, 19, 20, 21]. The effects induced by contractile density $\rho_0$ and chemo-mechanical coupling $\beta$ are likewise antithetical. Higher contractile densities would provide larger sources for ATP consumption and larger numbers of contractile dipoles per unit time, hence larger binding compared to unbinding rates, yielding more elongation and invasiveness of the cell [9]. Higher chemo-mechanical coupling, however, preserves chemo-mechanical energy transformation by inhibiting ATP consumption and non-conservative energy dissipation which, in turn, would preserve non-invasive cell shape [9].

### 3 Model Parametrization and Calibration

#### 3.1 Asymptotic analysis and allowable parameter ranges

Several mathematical constraints would restrict the ranges of allowable values for the model parameters. Before the steps are set forth, note that, as formerly confirmed in Sec. 2.2.2, the lower-branch solution would predominantly predict a spherical cell shape. Hence, the same parameter ranges derived herein for the special case of a spherical cell inside an infinite matrix can be later applied to general 3D cell-ECM domains with elongated (spheroidal) cell shapes.

By primitive definition, model tuning parameters $p$ and $c$ should adopt ranges that guarantee the trizonal solution. Accordingly, $0 < c \leqslant 1$ and $p > p_{\text{inf}}$, where the infimum power $p_{\text{inf}}$ is determined by the

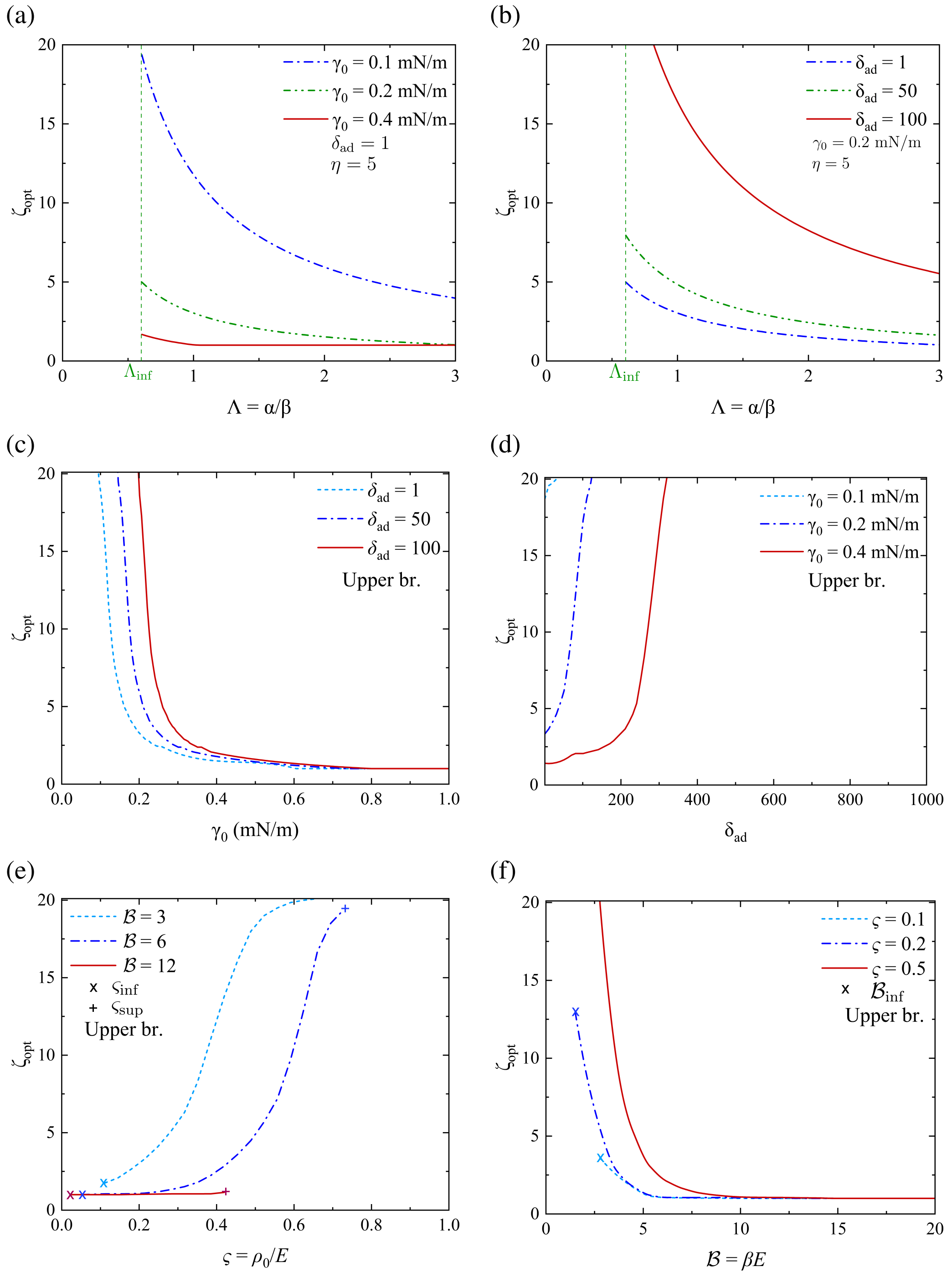


Figure S-6: **Effect of model parameters on highly motile optimal cell shape.** (a,b) Upper-branch (highly motile) optimum cell body aspect ratio at a sufficiently high ECM to cell stiffness ratio of $\eta \equiv E_{\mathrm{M}}/E = 5$ in terms of normalized polarization parameter $\Lambda \equiv \alpha/\beta$, with $(\varsigma, \mathcal{B}) = (0.2, 4.5)$: (a) at varying membrane–cortical tensile stiffness $\gamma_0$ (with $\delta_{\mathrm{ad}} = 1$), (b) at varying integrin contribution $\delta_{\mathrm{ad}}$ (with $\gamma_0 = 0.2$ mN/m); (c,d) in terms of $\gamma_0$ (with$\delta_{\mathrm{ad}} = 1$) and $\delta_{\mathrm{ad}}$ (with $\gamma_0 = 0.2$ mN/m), respectively; (e,f) in terms of normalized contractility density $\varsigma \equiv \rho_0/E$ (at varying $\mathcal{B} \equiv \beta E$) and normalized chemo-mechanical coupling $\mathcal{B} \equiv \beta E$ (at varying $\varsigma \equiv \rho_0/E$), with $\Lambda = 0.75$. Other cell parameters are identified by $E = 1.5$ kPa, $\nu = 0.27$. Adhesion binding affinity stiffness is fixed at $\gamma_1 = 0.001$ mN/m. Model tuning parameters are also fixed at $(p, c) = (6, 1/3)$ (a–d) or $(p, c) = (8, 1/3)$ (e,f).

nonnegativity condition of discriminant $\Delta$. That is

$$\Delta > 0 \;\therefore\; p > p_{\text{inf}} = \left[1 + 2\mathcal{B}'\varsigma\left(1 - \sqrt{1 + \frac{1}{\mathcal{B}'\varsigma'}}\right)\right]^{-1} \tag{S-32}$$

where $\mathcal{B}' = \beta' E$ (with $\beta'$ defined in advance) and $\varsigma' = \frac{1}{c}\varsigma$ with $\varsigma = \rho_0/E$. While Eq. (S-32) sets a lower limit to exponent $p$, it should be maintained well above the upper bound to this infimum at high levels of contractile density and/or chemo-mechanical coupling. The latter upper bound, as shown in Fig's S-7(e,f), is mathematically set by the $\mathcal{B}'\varsigma$ approaching infinity. Naming this product $\chi = \mathcal{B}'\varsigma = \beta'\rho_0$, one can obtain the supremum to $p_{\text{inf}}$ from its asymptotic value at $\chi \longrightarrow \infty$. To this end, the emerging $\infty - \infty$ ambiguity can be resolved by replacement of function $f(\chi) = \sqrt{1 + \frac{c}{\chi}}$ by its Taylor series, with $1/\chi$ transformed into a new variable such as $t$, thus $f(t) = \sqrt{1 + ct}$. Namely

$$\begin{aligned} \lim_{\chi\to\infty} f(\chi) &\equiv \lim_{t\to 0} f(t) = \lim_{t\to 0}\left(f(0) + f'(0)t\right) \\ &= \lim_{t\to 0}\left(1 + \frac{c}{2}t\right) = \lim_{\chi\to\infty}\left(1 + \frac{c}{2\chi}\right) \end{aligned} \tag{S-33}$$

which delivers the following asymptotic limit to $p_{\text{inf}}$:

$$\sup(p_{\text{inf}}) = \lim_{\chi\to\infty}\left[1 + 2\chi\left(1 - f(\chi)\right)\right]^{-1} = \frac{1}{1-c} \tag{S-34}$$

that entails adopting powers beyond this limit, *i.e.* $p > \frac{1}{1-c}$. For $c = 1/3$, for instance, the above supremum turns out 1.5 as shown in Fig's S-7(e,f).

Next, we evaluate the allowable range for the stress polarization parameter $\alpha$, in its normalized form $\Lambda = \alpha/\beta$. The general condition for this parameter is set by nonnegativity of $\eta_{\text{c}}$ in Eq. (S-31), which requires

$$\Lambda' > \frac{1}{p{\kappa_c}^{1-1/p}\left(1 - c\kappa_c\right)^{1+1/p}} \tag{S-35}$$

where $\Lambda' = \alpha/\beta'$. The lower limit to $\Lambda'$, however, can be evaluated from the minimum discriminant, *i.e.* $\Delta = 0$, which is determined by $\kappa_{\text{c}}^{(1)} = \kappa_{\text{c}}^{(2)} = q'/2$. Consequently, the infimum value of $\Lambda' = \alpha/\beta'$ would read

$$\Lambda'_{\text{inf}} = \frac{1}{p}\left(\frac{q'}{2}\right)^{1/p-1}\left(1 - \frac{q}{2}\right)^{-1/p-1} \tag{S-36}$$

where $q$ and $q'$ were defined in advance.

Figure S-8 depicts infimum and allowable ranges of $\Lambda = \alpha/\beta$ in terms of exponent $p$, normalized contractile density $\varsigma = \rho_0/E$ and normalized chemo-mechanical coupling parameter $\mathcal{B} = \beta E$. Since increasing exponent $p$ as well as decreasing normalized contractile density $\varsigma$ and chemo-mechanical coupling parameter $\mathcal{B}$ would lower the upper-branch predicted motility level, only a choice of higher polarization parameter (reflected in normalized $\Lambda$) would make amends for the reduced cell contractility as demonstrated by Fig. S-8.

Last, the infima (lower bounds) to both the normalized contractile density $\varsigma$ and chemo-mechanical coupling parameter $\mathcal{B}$ are set by the condition of nonnegativity of $\eta_{\text{c}}^{(2)}$ in Eq. (S-31), due to its emanation from $\kappa_{\text{c}}^{(2)}$, proving always lower than $\kappa_{\text{c}}^{(1)}$. Accordingly, the infimum $\varsigma$ and $\mathcal{B}$ would be determined by letting the denominator of $\eta_{\text{c}}^{(2)}$ vanish. That is, the $\kappa_{\text{c}}^{(2)}$ solution to

$${\kappa_{\text{c}}}^{q}\left(1 - c\kappa_{\text{c}}\right)^{2-q} = \frac{1}{p\Lambda'} \tag{S-37}$$

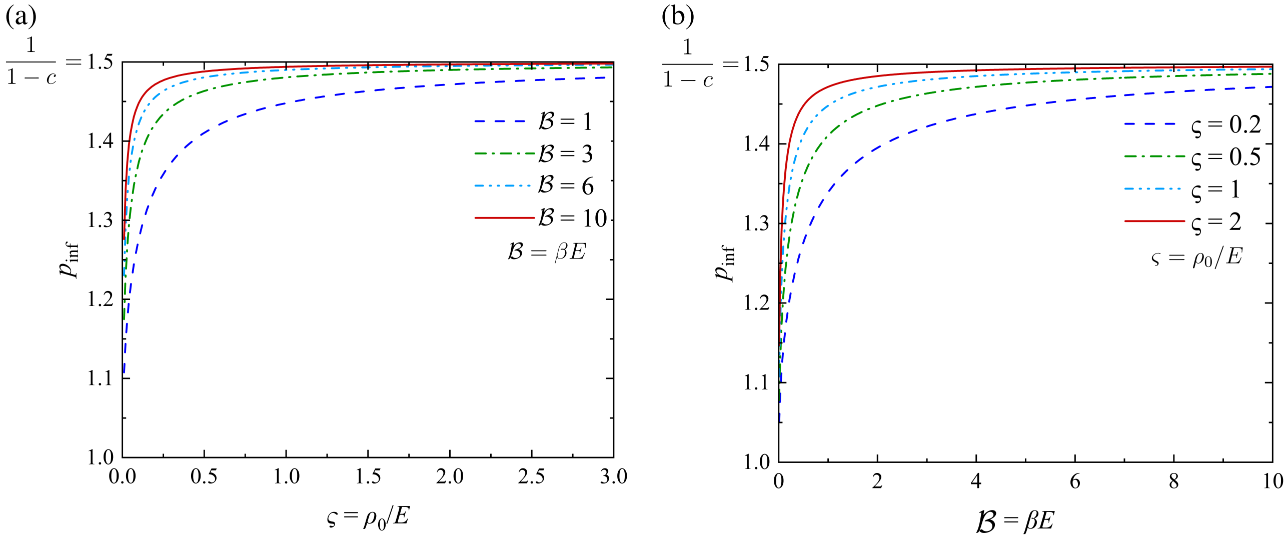


Figure S-7: **Allowable range of model tuning power $p$.** Infimum values of model tuning power $p$: (a) in terms of normalized contractility density $\varsigma \equiv \rho_0/E$, (b) in terms of normalized chemo-mechanical coupling parameter $\mathcal{B} \equiv \beta E$. Cell parameters are identified by $E = 1.5$ kPa, $\nu = 0.27$, $(\varsigma, \mathcal{B}, \Lambda) = (0.2, 4.5, 1)$.

would yield $\kappa^{(2)}_{\text{cmin}}$. On the other hand, from the $2\kappa^{(2)}_{\text{c}} = q' - \sqrt{\Delta}$ correlation in Eq. (S-31), one can easily verify the following infimum to the $\varsigma'\mathcal{B}'$ product:

$$(\varsigma'\mathcal{B}')_{\text{inf}} = \left(q' - \kappa^{(2)}_{\text{cmin}}\right) p\, \kappa^{(2)}_{\text{cmin}} \tag{S-38}$$

Note further that the sign of $\eta_{\text{c}}$ in Eq. (S-31) is partly affected by the sign of $\mathcal{G}$. Hence, the positivity of $\mathcal{G}$ imparts an additional constraint on $\mathcal{B}$. In the case of a cell embedded in an infinite ECM, such constraint requires that $\mathcal{B} > \frac{1}{1-2\nu}$. Hence, infimum $\varsigma$ and $\mathcal{B}$ would be obtainable from the combination of Eq's (S-38) and (S-37) with the above additional constraint, which delivers

$$\begin{cases} \forall \mathcal{B}: \quad \varsigma_{\text{inf}} = \dfrac{\left(q' - \kappa^{(2)}_{\text{cmin}}\right) pc\, \kappa^{(2)}_{\text{cmin}}}{\mathcal{B}'} \\[2ex] \forall \varsigma: \quad \mathcal{B}'_{\text{inf}} = \min\left\{ \dfrac{\left(q' - \kappa^{(2)}_{\text{cmin}}\right) pc\, \kappa^{(2)}_{\text{cmin}}}{\varsigma'} \,,\, \dfrac{1}{\Delta G(1-2\nu)} \right\} \end{cases} \tag{S-39}$$

All the same, the maximum allowable values for $\varsigma$ and $\mathcal{B}$ are derivable from the minimum value of the discriminant in Eq. (S-31), *i.e.* $\Delta = 0$. Therefore, the supremum (upper bound) value for $\varsigma$ and $\mathcal{B}$ would be obtained from the infimum value of the other. That is

$$\begin{cases} \forall \mathcal{B}: \quad \varsigma_{\max} = \frac{p}{4c} \frac{q^2}{\mathcal{B}'} \;\therefore\; \varsigma_{\sup} = \frac{p}{4c} \frac{q^2}{\mathcal{B}'_{\text{inf}}} \\[2ex] \forall \varsigma: \quad \mathcal{B}'_{\max} = \frac{p}{4c} \frac{q^2}{\varsigma} \;\therefore\; \mathcal{B}'_{\sup} = \frac{p}{4c} \frac{q^2}{\varsigma_{\text{inf}}} \end{cases} \tag{S-40}$$

where $\mathcal{B}'_{\text{inf}}$ and $\varsigma_{\text{inf}}$ follow Eq. (S-39).

From Eq's (S-38)–(S-40), the dimensionless parameters $\mathcal{B}' = \beta' E$ and $\varsigma = \rho_0/E$ are found to be physical conjugates. That is, at a given steady state, one should decrease with the other. Note that, unlike $\varsigma'$

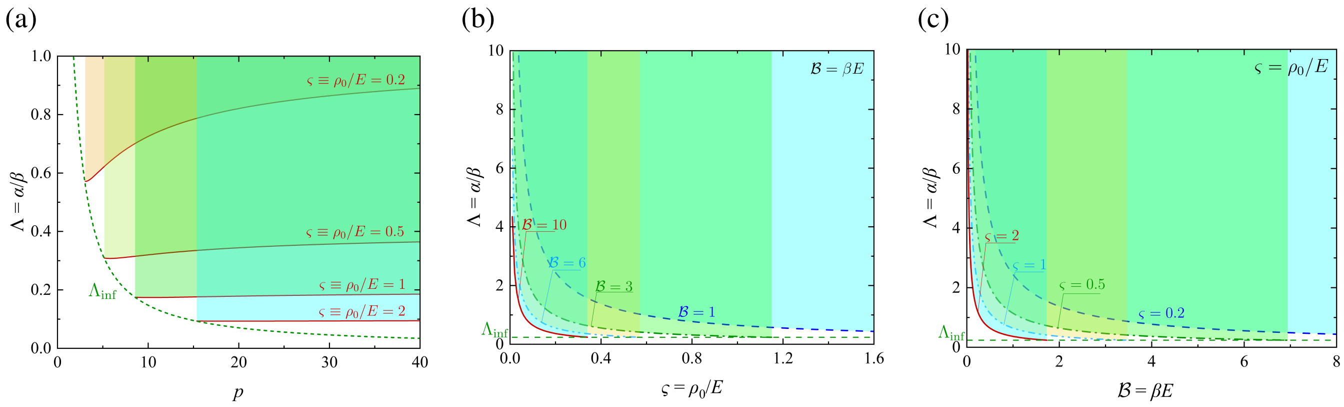


Figure S-8: **Allowable range of stress polarization parameter.** Ranges of the normalized polarization parameter $\Lambda \equiv \alpha/\beta$ with its infimum value showing as lower bound: (a) vs. model tuning power $p$, (b) vs. normalized contractility density $\varsigma \equiv \rho_0/E$ (at select normalized chemo-mechanical coupling parameters $\mathcal{B} \equiv \beta E$), (c) vs. normalized chemo-mechanical coupling parameter $\mathcal{B} \equiv \beta E$ (at select normalized contractility densities $\varsigma \equiv \rho_0/E$).

and $\varsigma$ that would differ in a $c$ factor, $\mathcal{B}'$ and $\mathcal{B}$ can be considered equal without loss of generality inasmuch as $\Delta G$, as primitively defined in Eq. (S-15), is merely a scaling factor that can easily get absorbed into $\beta$. Figure S-9 shows example curves for allowable ranges of $\mathcal{B}$ and $\varsigma$ at various normalized polarization parameters $\Lambda$, each plotted vs. the other.

Once the above-discussed model parameters are defined, the bounds to trizonal solution can be evaluated in terms of the minimum and maximum critical stiffness ratios $\eta_c^{(1)}$ and $\eta_c^{(2)}$, as indicated by Fig. S-10. The lower value $\eta_c^{(1)}$ would then represent a measure of the minimum ECM stiffness that triggers shape transition, cell elongation and metastatic invasiveness. Besides, the upper value $\eta_c^{(2)}$ signifies the maximum ECM stiffness which generates two possible steady states of the cell, one non-invasive and one metastatic. Any value beyond $\eta_c^{(2)}$ would represent a fully invasive cell state. Nevertheless, cell elongation and invasiveness in such high ranges of ECM stiffness would decrease with increasing stiffness due to the highly dense, intertwined (basket-weave) network of fibers, as earlier elaborated on in Sec. 2.2.2.

#### 3.2 Model calibration against experiment

The sequel provides the grounds for model calibration against experimental histograms through the parametric sweep mode introduced in Eq. [6] of the manuscript.

##### 3.2.1 *Parametric sweep*

The calibration algorithm primarily invokes a full-spectrum data library calculated from model predictions with varying contractile density according to the following variation law:

$$\bar{\rho_0} = \rho_0(1 + g_0\phi) \qquad \text{(S-41)}$$

which is a replica of Eq. [8] in the manuscript. Here, $g_0$ denotes an intensity coefficient obeying $0 < g_0 < 1$ (fixed at every case and varying from case to case) and $-1 \leqslant \phi \leqslant 1$ is the main sweeping parameter. While $g_0$ sets the variation range of contractile density, its core value $\rho_0$ also varies such that $0.1 \leqslant \varsigma \equiv \rho_0/E \leqslant 0.6$. Besides varying $g_0$ and $\varsigma$, the parametric library takes into account varying membrane–cortical tensile stiffness $\gamma_0$, integrin contribution $\delta_{\rm ad}$, and normalized stress polarization parameter $\Lambda$. Example plots with

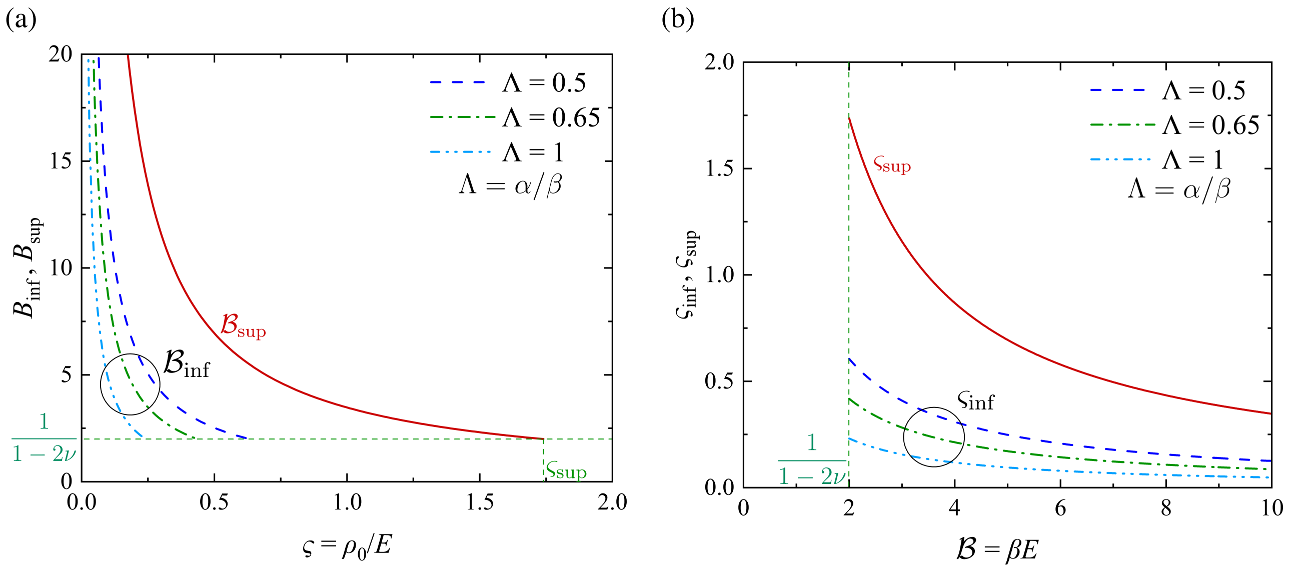


Figure S-9: **Allowable ranges of contractile density and chemo-mechanical coupling.** Infimum and supremum values of normalized contractile density $\varsigma \equiv \rho_0/E$ and chemo-mechanical coupling parameter $\mathcal{B} \equiv \beta E$ at select values of normalized polarization parameters $\Lambda \equiv \alpha/\beta$. Cell parameters are identified by $E = 1.5$ kPa and $\nu = 0.27$. Model tuning parameters are also fixed at $(p, c) = (6, 1/3)$.

above variations vs. $\phi$ is shown in Fig. S-11 so as to include the optimum cell body aspect ratios lying within the experimentally observed [1,5] range. The values result from the same analysis procedure laid out for the linear model. Each point on the referred plots in Fig. S-11 was obtained from simultaneous solution of equilibrium and constitutive equations scanned throughout a prescribed range of cell body aspect ratios $1 \leqslant \zeta \leqslant 20$. Energy differences $\Delta U_{\text{tot}}$ were then exported to a post-processing subroutine to evaluate the optimum aspect ratio $\zeta_{\text{opt}}$ from the local minimum to the $\Delta U_{\text{tot}}$ vs. $\zeta$ curve.

##### *3.2.2 Multimodal histogram generation*

This subroutine would generate the best histograms provided the first step was paid to predict the best parameter set ($(\varsigma, g_0)$, $(\Lambda, \mathcal{B})$ and $(\delta_{\text{ad}}, \gamma_0)$) associated with the least squared difference acquired between the model predictions and the main data peaks on the histogram of interest. Thereupon, the entire histographic $\zeta$ distribution was simulated by application of a replicated multimodal probability distribution subroutine recently built in MATLAB as the *Gaussian mixture model* (`fitgmdist`) [34]. The latter utilized the iterative *Expectation Maximization* (EM) algorithm which operates mainly as function of the data peaks (at least the largest ones) and the number of desired peaks, *alias* modes ($n_{\text{p}}$). The above algorithm proceeds through the following steps [34]:

1. For each data realization, the algorithm computes posterior probabilities of component memberships in the form of a $N$-by-$n_{\text{p}}$ matrix ($N$ denoting data count), whose $ij^{\text{th}}$ element is provided by the posterior probability that the observed $i^{\text{th}}$ data is from component $j$.

2. The component-membership posterior probabilities at hand are regarded as weights, upon which the algorithm estimates each and every component's mean, covariance matrix and mixing proportion by application of the maximum likelihood.

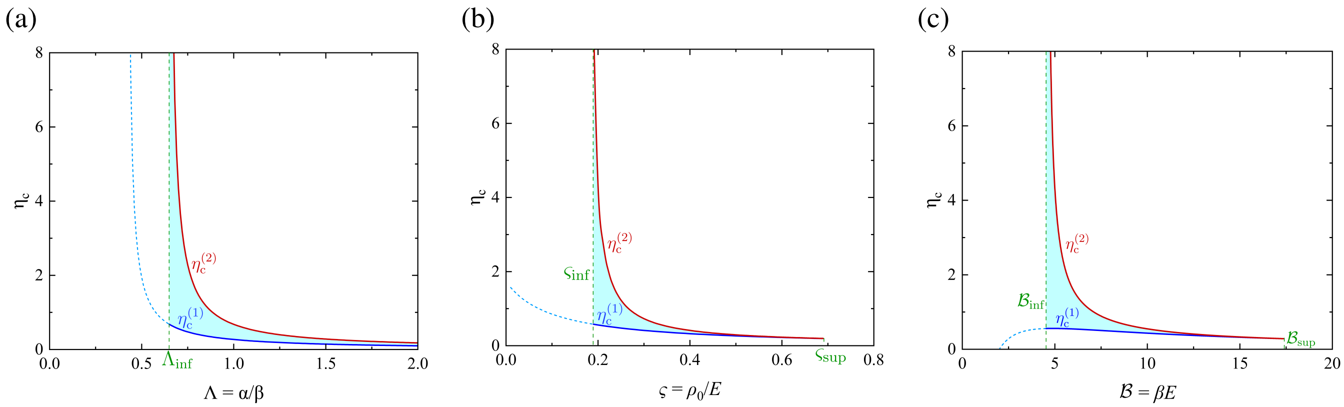


Figure S-10: **Minimum and maximum critical stiffnesses for trizonal cell state.** Bounds to the stiffness overlapping interval $\eta_{\rm c}^{(1)}$ and $\eta_{\rm c}^{(2)}$: (a) in terms of the normalized polarization parameter $\Lambda \equiv \alpha/\beta$ (with $(\varsigma, \mathcal{B}) = (0.2, 4.5)$), (b) in terms of normalized contractile density $\varsigma \equiv \rho_0/E$ (with $\mathcal{B} = 4.5$), (c) in terms of the normalized chemo-mechanical coupling parameter $\mathcal{B} \equiv \beta E$ (with $\varsigma = 0.2$). Other cell parameters are identified by $E = 1.5$ kPa and $\nu = 0.27$. Model tuning parameters are also fixed at $(p, c) = (6, 1/3)$.

3. The above two steps are iterated until a prescribed or default-valued convergence is achieved.

**Choice of Initial Conditions:** Inasmuch as the resulting local optimum from the EM algorithm might depend on the initial conditions, `fitgmdist` can be equipped with several options for the most judicious choice of initial conditions, including random component assignments for data realizations and the so-called *k-means*++ algorithm [35, 36]. The latter algorithm uses heurism to find centroid seeds for *k-means* clustering. To this end, the algorithm assumes the number of clusters as being $n_{\rm p}$, thereby choosing the initial parameter set as follows:

1. Selecting the component mixture probability as a uniform probability given by $p_i = \frac{1}{n_{\rm p}}$, with $i = 1, 2, ..., n_{\rm p}$.

2. Selecting identical diagonal covariance matrices as $\sigma_i = \text{diag}(a_1, ..., a_{n_{\rm p}})$, with $a_j = \text{var}(X_j)$ (denoting variance).

3. Selecting a uniform component center equal to its first initial value $\mu_1$ from all data points in the data set $\boldsymbol{X}$.

4. Selecting the $j^{\rm th}$ centroid as:

   – Computing the Mahalanobis distances from each observation to each centroid, and assigning each observation to its closest centroid.

   – Computing the $j^{\rm th}$ centroid from data set $\boldsymbol{X}$ with a probability being proportional to the distance from itself to the closest center, given by

$$p_{cj} = \frac{d^2(X_m, \mu_k)}{\sum_{h, X_h \in M_k} d^2(X_h, \mu_k)} \tag{S-42}$$

   where $m = 1, ..., N$, $k = 1, ..., j-1$, with $d(X_m, \mu_k)$ denoting the distance between observations $m$ and $\mu_k$, and $M_k$ denoting the set of all observations closest to centroid $\mu_k$ while $X_m$ belongs to $M_k$.

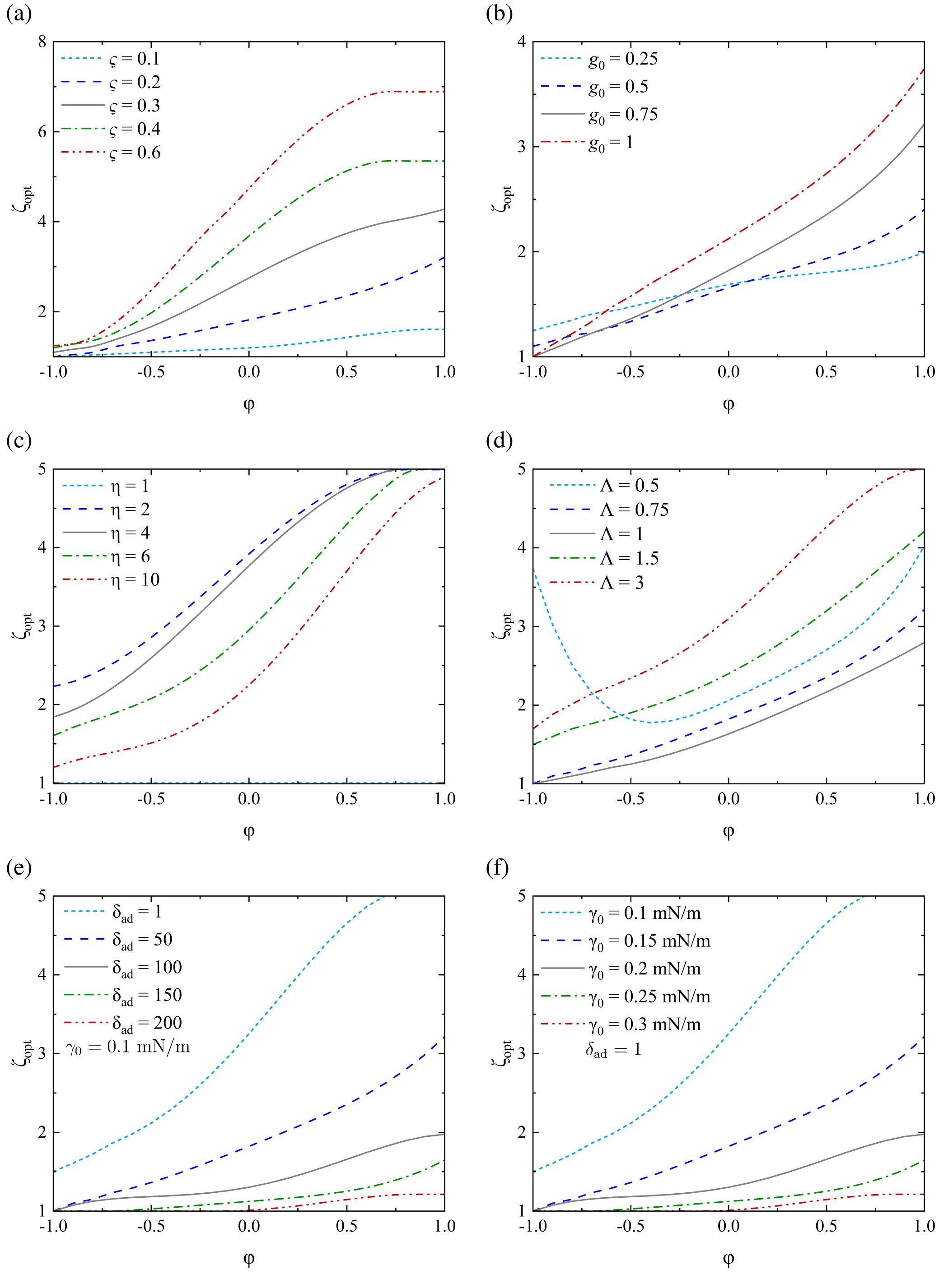


Figure S-11: **Parametric contours of contractile density.** Variation of contractile density $\bar{\rho}_0$ as perturbed from a core value $\rho_0$ through the sweeping parameter $\phi$: (a) at fixed stiffness ratio of $\eta = 3.5$ and select core values of $\rho_0$, with $g_0 = 0.75$, $\mathcal{B} \equiv \beta E = 5$, $\Lambda \equiv \alpha/\beta = 1$ and interfacial parameters $\gamma_0 = 0.15$ mN/m, $\delta_{\text{ad}} = 1$; (b) at select intensity parameters $g_0$, with the same abovementioned parameters (excluding fixed $g_0$) and $\varsigma = 0.2$; (c) at select stiffness ratios and the same abovementioned parameters; (d) at select normalized polarization parameters $\Lambda$, fixed stiffness ratio of $\eta = 3.5$ and the same abovementioned parameters; (e) at select integrin contributions $\delta_{\text{ad}}$, fixed values of $\gamma_0 = 0.1$ mN/m and $\Lambda = 0.75$, and the remaining parameters equaling the abovementioned; (f) at select membrane–cortical tensile stiffnesses $\gamma_0$, fixed values of $\delta_{\text{ad}} = 1$ mN/m and $\Lambda = 0.75$, and the remaining parameters equaling the abovementioned. Adhesion binding affinity stiffness and model tuning parameters are also fixed at $\gamma_1 = 0.001$ and $(p, c) = (10, 0.5)$, respectively.

5. Repeating step 4 until $n_p$ centroids are selected.

For more information, refer to [35] and MATLAB's user manual.

Note that, at least within the range of data at hand, bimodal distribution ($n_p = 2$) was found to normally generate the best coincidence with experimental histograms, and unimodal distribution was found optimal only in the few cases with a large peak and monotonic data distribution apart from the peak. No trimodal or higher-order distribution was found optimal. Note also that the parameter sets for nearly spherical cells ($1 \leqslant \zeta < 1.2$) were extracted from the model's lower branch and higher ratios were evaluated using the upper branch.
